# Social Isolation Alters Hippocampal miR-30e-5p Expression and Impairs Pattern Separation–Related Behaviour in Adult Mice

**DOI:** 10.64898/2026.06.24.734185

**Authors:** Amina McDiarmid, Amanda Kiemes, Gargi Mandal, Sandrine Thuret, Cathy Fernandes

**Author notes:** S.T. and C.F. equally contributed.

## Abstract

Social isolation is commonly used to model social stress and is a known risk factor for depression, with impacts on hippocampal function and postnatal neurogenesis. However, most studies focus on social isolation in juvenile mice isolation during adolescence, leaving the effects of prolonged adult isolation less understood. Post-transcriptional regulation of gene expression by microRNAs (miRNAs) plays a role in hippocampal function, and altered miRNA, as well as gene expression, has been reported in the hippocampus of mice exposed to social isolation. A single-nucleotide polymorphism in miR-30e in humans is associated with increased expression of the mature miRNA, impaired cognition, electroencephalogram waveform latency, depression, and schizophrenia. We investigated whether adult isolation in mice alters gene regulation via microRNAs, particularly miR-30e-5p, and affects hippocampal function. In adult BALB/c male mice, 10 weeks of isolation increased miR-30e-5p expression in the ventral hippocampus, reduced its target gene *Neurod1*, and impaired hippocampal-dependent cognition (object pattern separation), without clear anxiety- or depression-like behaviours. Isolated mice also showed a blunted response to acute stress. These findings suggest that adult social isolation affects hippocampal function through post-transcriptional gene regulation, highlighting a role for miR-30e-5p in neurogenesis and cognition in response to psychological stress.

## 4 Introduction

Persistent feelings of low mood and anhedonia are the two major clinical hallmarks of depression (American Pyschiatric Association and Gunderson, 2013). Symptoms of depression often occur along with anxiety suggesting the conditions are linked, perhaps via partly overlapping aetiology (Cloninger, 1990; Watson et al., 1995). Interconnected environmental and genetic risk factors associated with cognitive, emotional, psychological and biological disturbances could constitutively interact with each other to precipitate symptoms of MDD and/or anxiety (Grotzinger et al., 2026). Social stress models depression-and anxiety-like behaviours *in vivo*, and social isolation is particularly important since it is consistently associated with anxiety and depression in humans.

Social withdrawal is a common behaviour in anxiety and depression, and social isolation a common risk factor for both (Kim et al., 2025; Wilkialis et al., 2021; Zhu et al., 2024). Early work demonstrated that mice are sensitive to altered social environment. Individual housing in mice induces behavioural changes compared to standard laboratory conditions of group or paired housing (Valzelli, 1973). Thus, the model has been widely adopted to assess the impact of a key psychological stress in affective disorders, social isolation. Individually housed mice display behaviours analogous to anhedonia and despair (Atrooz et al., 2021; Cryan and Holmes, 2005; Scheggi et al., 2018). Affective-related behaviours and spatial cognition also have previously been reported in adult mice that were exposed to early life social isolation starting during adolescence (Hall et al., 1998; Hanif et al., 2025). Social isolation of rodents in early adulthood or old-age mimics aspects of depression and anxiety (Aguiar et al., 2014; Magalhães et al., 2024; Mamedova et al., 2024), impairs spatial memory (Zorzo et al., 2019) and stress response (Serra et al., 2005). However, the neurobiological processes, psychological mechanisms and molecular pathways affected by social isolation remain unclear.

The hippocampus plays a critical role in contextual and episodic memory encoding/retrieval, which subsequently impacts negative bias of information processing (Chen et al., 2019; Hamilton and Gotlib, 2008; Radzicki et al., 2025). Should a negative or stressful experience occur in a context similar to a previously encoded benign memory, the ability to recall events separately (despite contextual similarity) is crucial to prevent catastrophic interference between benign and negative memories and, in so doing, to avoid negative psychological biases (Egeland et al., 2015). Associative memory, which is the recall of memories from partial cues and older memories during novel experiences, requires accurate encoding and recall of prior memories in order to distinguish it from the newly encoded memories of the present experience (Duncan et al., 2014). Such type of memory processing is characteristic of the hippocampus and place a huge demand on pattern separation. Pattern separation is a process by which memories of contextually similar experiences undergo separate neural encoding to form non-overlapping representations of highly similar information, allowing the brain to distinguish between similar events and prevent memory interference (Ngo et al, 2021). Pattern separation could influence mood by preventing overgeneralisation, a psychological feature of depression (Bakker et al., 2008; Shelton and Kirwan, 2013) and anxiety (Kheirbek et al., 2012). Thus, there is a developing role for this form of cognition in affective disorders such as depression and in anxiety.

Pattern separation has been shown to be associated with hippocampal activation using functional magnetic resonance imaging (Kirwan and Stark, 2007). Computational modelling of hippocampal neural networks suggests the dentate gyrus (DG) granule neurons are especially able to separate similar incoming projections from the entorhinal cortex (EC) before relaying this information to CA3 pyramidal cells via the mossy fibres (Hu et al., 2025; Norman and O’Reilly, 2003). Indeed, selective lesions of the DG disrupt pattern separation in rats (Morris et al., 2012).

The DG is one of two identified regions in the mammalian brain where neurogenesis, the process by which new neurons are persistently produced from a pool of proliferative neural progenitor cells, occurs into adulthood and old age in humans (Dumitru et al., 2025; Spalding et al., 2013) and many other mammalian species including mice. Both acute and chronic stress reduce hippocampal neurogenesis in a dose-dependent manner (Kempermann, 2002; Schoenfeld et al., 2017). In particular, social isolation reduced neurogenesis in the rodent DG and increased depressive-/anxiety-like symptoms (Du Preez et al., 2021; Lieberwirth et al., 2012; Zlatković et al., 2014). Conversely, pro-neurogenic behaviours, such as running has been associated with increased pattern separation (Creer et al., 2010; Hvoslef-Eide and Oomen, 2016). Pattern separation is dependent on adult hippocampal neurogenesis, since abating it in mice impairs discrimination between overlapping spatial locations (Clelland et al., 2009). Furthermore, evidence suggests dorsal and ventral hippocampus in rodents, analogous to the posterior and anterior in humans, have distinct spatial and functional substructures (Fanselow and Dong, 2010; Moser and Moser, 1998), with differential gene expression profiles (Beletskiy et al., 2022; Lee et al., 2017). Although the functional mechanisms of adult-born neurons in the ventral hippocampus are critical in the behavioural response to social stress (Anacker et al., 2018), the underlying molecular mechanisms remain elusive.

Of particular interest in the context of molecular regulation in hippocampal cognition and mood is microRNA-30e-5p (miR-30e-5p). A single-nucleotide polymorphism (SNP) in its precursor (pre-miR-30e) is associated with increased levels of mature miR-30e-5p and has been linked to major depressive disorder (MDD) and schizophrenia (SCZ) (Xu et al., 2010a, 2010b). This variant is also associated with increased p300 latency in EEG recordings during oddball paradigm tasks in MDD (Xu et al., 2010b). The oddball paradigm tests the subjects’ ability to distinguish between previously experienced and novel stimuli, and such event-related potentials are widely used as a parameter of mnemonic information processing and spatial separation (Chot et al., 2020; Polich, 2007; Warchoł and Zając-Lamparska, 2023). Outcomes are measured using EEG and event-related potentials such as p300 latency (Sutton et al., 1965) are widely used as a parameter of mnemonic information processing. Indeed, the hippocampus (and its connections to other pivotal brain regions) is believed to make a major contribution to p300 EEG recordings (Ludowig et al., 2010). Elevated miR-30e-5p expression in rodents has been linked to impairments in spatial memory (Xu et al., 2016, 2015). Abnormal event-related potentials signify impaired mnemonic binding which is detrimental to processing of the contextual environment (Guillem et al., 2003). As pattern separation is an artefact of mnemonic processing, taken together this evidence suggests that increased miR-30e-5p may contribute to impaired pattern separation-like cognition and broader hippocampal dysfunction but the downstream mechanisms require further elucidation.

Social isolation disproportionately affects males (Röhr et al., 2022), in whom clinical (Chi et al., 2024; Vallée, 2025) and biological (Umberson et al., 2022) responses to social isolation in significantly worse relative to females. In this study a model of social isolation in adult male mice was used to investigate the possible regulatory interaction between miR-30e-5p and putative downstream target mRNAs. We hypothesised that 10 weeks of individual housing would impair object pattern separation, induce anxiety- and depression-like behaviours and modify circulating corticosterone levels. We further predicted, the same social isolation paradigm run in behaviour test-naïve mice would alter hippocampal miR-30e-5p expression based on prior evidence linking miR-30e-5p overexpression to impaired hippocampal cognition and alter expression of miR-30e-5p target genes identified from *in silico* database mining. We found behavioural measures, object pattern separation, post-stress serum concentration of CORT, expression of miR-30e-5p and a predicted target relevant to hippocampal neurogenesis (*Neurod1)* are altered in male BALB/c mice after 10 weeks of social isolation during adulthood.

## 5 Materials and methods

### 5.1 Animals

Forty, adult (7-8 week-old) male BALB/cAnNCrl (BALB/c) inbred mice were purchased from Charles River Laboratories (Margate, UK). All animals were maintained on a 12-hour light/dark cycle (lights on 0800) in a temperature-controlled room (19-22°C) at 55% humidity in the Biological Services Facility (BSF) at the Institute of Psychiatry, Psychology and Neuroscience (King’s College London, Denmark Hill Campus).

### 5.2 Modelling social isolation by individual housing

Male mice were matched for body weight (g) on delivery and then randomly assigned to one of two housing conditions. Males were housed either in pairs (PH) or single-housed (social isolated group, SI). PH males were housed in a large cage (60cm x 30cm x 20cm: Techniplast, London, UK) (n=20/housing group). PH males were provided with additional paper nesting material (sufficient for both mice) with an additional grey polycarbonate tunnel and red triangular Techniplast Mouse House^©^ which has two access points (one at cage base level and one leading from cage base to the roof of the house), a roof acting as a shelf or platform for climbing and a large nesting/shelter area or individually (SI) in a standard mouse cage (36cm x 20cm x 14cm: Techniplast, London, UK) with a cardboard house (Shepherd Shack, LBS Biotech, Horley, UK) and paper nesting material (Datesand Ltd, Manchester, UK). Larger cages were used for pair-housed mice for welfare reasons since BALB/c males naturally show territorial behaviour and nest building is an important natural behaviour within this strain (Gaskill et al., 2013; Lee and Wong, 1970). Therefore according to guidance from NC3Rs, sufficient nesting/housing materials within cages to allow socially housed cagemates to create individual nests and territories for welfare reasons. Cages included a 1cm layer of sawdust bedding (Litaspen premium, Datesand Ltd, Manchester, UK) and cleaning was carried out fortnightly. These housing conditions were maintained for a total of 10 weeks. Food (5053 – PicoLab Rodent Diet 20, Lab Diet, St. Louis, USA) and water (available in two bottles per cage) were placed in the metal grid cage lid and available *ad libitum*.

### 5.3 Ethics

All housing, handling and experimental procedures were performed in compliance with the local ethical review panel of King’s College London under a UK Home Office project license (PPL: 70/7506) granted in accordance with the Animals (Scientific Procedures) Act 2012.

### 5.4 Body Weight

A baseline measure of body weight (g) was taken upon arrival (designated day 0) and weekly thereafter (in the morning on day 7, 14, 21, 28, 35, 42, 49, 56 and 63).

### 5.5 Food and water consumption

A 24 hr measure of food consumption was recorded weekly. To determine the amount of food eaten over a 24-hour period, the amount of food (g) available on the first day was taken as a baseline with a follow-up measurement taken 24 hours later. Similarly, water consumption (g) was also measured over a 24h period to determine daily water intake with control for spillage determined from 6 control cages with the same cage enrichment, placed on the cage racks but not holding any mice (referred to as spill cages).

### 5.6 Behavioural Assessment

A battery of behavioural assessments was performed (Supplementary Methods). Repeated measures of coat state (weeks 0-10) and sucrose preference (weeks 0, 2, 4, 6 and 10) were performed, with one-off assessments of open field behaviour (week 6), olfaction (week 7), social dominance (week 8), object pattern separation (week 8), splash test (week 9), novelty supressed feeding test (week 10) and concluding with Porsolt swim test (week 10) (Supplementary Figure 1 and 2). Procedures were carried out in half of the mice (n=10 individually housed and n=10 pair-housed) retaining the other half test-naïve for gene expression analysis. Cages were not changed or cleaned within 24h of, or on the day of, testing. All behavioural testing was performed during the light phase of the light: dark cycle between 09:00 and 17:00. During testing, all animals were placed in the start location within each test, and after each trial, boli and urine were removed from the test arena, which was subsequently cleaned with 1% Anistel® solution (Tristel Solutions Ltd, Suffolk, UK). Light levels were task specific (described in each test in subsequent subsections), but remained constant, within each test, for all experimental groups. All tests were recorded using a camera positioned above the test arena, and for some tests the animal’s movement was tracked using EthoVision software (Noldus Information Technologies, Wageningen, The Netherlands). At the end of testing, mice were returned to their home cage, and then to the housing room. An experimenter blind to the housing group of the mice performed any live or post-acquisition scoring of the behaviours.

### 5.7 Object Pattern Separation

The Object Pattern Separation (OPS) task was conducted using the open field arena as previously described (van Hagen et al., 2015). Briefly, the test is an adapted version of object recognition tasks commonly used in rodents to investigate cognition. In contrast to other object recognition tasks where rodents are exposed to novel versus familiar objects, or objects displaced across a large distance in an arena, this task determines if animals can detect an object displacement across a small distance (5-15cm). The Object Pattern Separation protocol consists of a habituation period and test trials (Supplementary Methods, Supplementary Figure 2G, 2H and 3).

### 5.8 Behaviour Scoring Validation

Behaviour was manually scored using MATLAB R2017a for the following tests: Porsolt swim test, OPS test and Splash test. To assess intra-rater reliability five trials from each test were scored five times. Inter-rater reliability for results generated manually post-acquisition from recorded footage of the Porsolt swim and OPS trials. Scoring was performed by two blinded experimenters and intra-/inter-rater reliability was calculated using the intra-class correlation coefficient (ICC) (Bartko and Carpenter, 1976), where the validation threshold is ICC> 0.80. Inter-rater and intra-rater reliability as determined by ICC was >0.99 for both the OPS test and Porsolt swim test.

### 5.9 Blood sample collection

Whole blood was collected by incision method from the lateral tail vein (Sadler and Bailey, 2013). In brief, ∼50 µl blood as collected drop wise in capillary tubes (potassium-EDTA microvette CB 300 tubes, Sarstedt, Nümbrecht, Germany) from a small incision made using a sterile blade (American Safety Razor Blade, Persona) on the underside of the tail ∼15mm from the base. Samples used in the current study were preserved on ice prior to separation by centrifugation to minimise any degradation of corticosterone. Blood samples were collected on 3 occasions in the 10^th^ week: 1) 24 hours prior to the Porsolt swim test (from 09:00 to 17:00), 2) 30 minutes post-Porsolt swim test (time-matched to the pre-swim baseline collection time for each animal) and 3) immediately prior to sacrifice (between 17:00-18:15 to minimize variation due to natural circadian fluctuations in corticosterone (Dalm et al., 2005). Plasma was extracted following centrifugation (12,000rpm for 10mins at 4°C).

### 5.10 Hippocampal tissue dissection and collection

After 10 weeks of isolation, animals were euthenised by cervical dislocation and decapitation. Brain samples were dissected and collected fresh into sterile, nuclease-free 1.5mL tubes (0030123328, Eppendorf, UK) and snap-frozen on dry ice. Dissections were carried out to isolate frontal cortex, cerebellum and the hippocampus. The hippocampus was further dissected into left ventral, left dorsal, right ventral and right dorsal hippocampi. To do this, the whole hippocampi were removed and dorsal/ventral and anterior/posterior orientation retained relative to the rest of the brain with the hippocampus laid flat on the medial side. The ventral hippocampus is identifiable by a small hump-like protrusion. To dissect the ventral from dorsal, the hippocampus was divided into 5 equal sections from top (dorsal) to bottom (ventral). The top two-fifths were dissected as one piece of tissue, while the bottom 2/5s were dissected as another with the central remaining 5 discarded. The top was taken as the dorsal hippocampus and the bottom as the ventral. Methods as previously described by (Kelly et al., 2010) and dissections validated using expression assay of gene *Pde11a4*, which enriched in the ventral relative to dorsal hippocampus.

### 5.11 Corticosterone Assay

Corticosterone (CORT) levels in plasma were assayed using the commercial corticosterone enzyme-linked immunosorbent Assay kit (ELISA) (ADI-900-097; Enzo Life Sciences AG; Lausen, Switzerland) as per the manufacturer’s instructions. For each plasma sample, 10μl was diluted 1:25 with steroid displacement reagent and assay buffer. CORT standard curve samples were prepared via serial dilution (starting concentration 200ng/μL) to generate a range of 5 concentrations (20, 4, 0.8, 0.16, and 0.032 ng/ml). ELISA plates were pre-coated with corticosterone antibody (donkey anti-sheep IgG). All samples were analysed in duplicate. The intra-assay variability ranged from 6.6% to 8.4% and sensitivity = ±26.99 pg/mL.

### 5.12 RNA extraction

Tissue samples were snap frozen in dry ice then stored at -80°C. Tissues were lysed and homogenized in TRIzol reagent (15596026, ThermoFisher) using sterile single-use disposable plastic pestles which were inserted into a hand-held, rotating mechanical homogenizer. RNA was then extracted using a chloroform phase separation (to produce an RNA-containing aqueous phase, and protein/DNA-containing organicand interphase) followed by isopropanol precipitation including a purification using sodium acetate (R1181, ThermoFisher) and 80% ethanol (VWR). Concentration ([RNA]ng/µL) and sample purity (260/280 and 230/260 ratio) were determined using a spectrophotometer (NanoDrop One, ThermoFisher). Samples with low purity were eliminated, re-extracted or purified and washed before downstream analysis.

### 5.13 Quantitative real-time PCR

For miRNA expression analysis, the TaqMan miRNA Reverse Transcription kit and TaqMan miRNA Assay was used as per manufacturer’s instructions. In brief, hairpin reverse transcription primers specific to mmu-miR-30e-5p were used to reverse transcribe target miRNAs and endogenous reference gene (small non-coding RNA U6) using MultiScribe enzyme (cycling conditions as follows: 30 mins at 16°C → 30 mins at 42°C→5mins at 85°C). Then TaqMan quantitative PCR (qPCR) using TaqMan Universal MasterMix II (no UNG) with target specific probes was used to generate data for the purpose of quantifying relative expression (cycling conditions as follows 10mins at 95°C → 40 cycles of 15 secs at 95°C + 60secs at 60°C). Fold-change relative to control samples, normalised to an endogenous reference gene, was calculated from TaqMan qPCR using the ΔΔC_T_ method due to TaqMan probes optimisation for 100% (±10%) efficiency.

For mRNA expression analysis, qPCR was carried out using HOT FIREPol® EvaGreen qPCR Mix Plus (ROX) (SolisBiodyne). Prior to this, extracted RNA was treated with DNase to remove DNA contamination (Ambion TURBO DNA-*free*^TM^ kit, AM1907). Reverse transcription (SuperScript III or IV Invitrogen, 18080093) was carried out according to instructions using random hexamer primers (Invitrogen, N8080127) to create a cDNA library. Primer design was conducted such that either the forward or reverse primer targeted an exon-exon boundary within the gene of interest for the purpose of targeting mRNA (and not genomic DNA). Exon-exon boundaries and their corresponding sequences were manually selected and downloaded from the National Center for Biotechnology Information (NCBI) Gene database (https://www.ncbi.nlm.nih.gov/gene). The PrimerQuest^TM^ online, open-access tool (Integrated DNA Technologies) was used to generate possible primer sequences and selected using Primer-BLAST (NCBI) based on specificity. Primer-BLAST was used to predict off-target amplification and only primers with no predicted off-target amplicons were selected. Primer oligonucleotide molecules were synthesised to order by Integrated DNA Technologies with primer sequences as follows; *Neurod1* forward 5’-ACCTTTTAACAACAGGAAGTGGA-3’, *Neurod1* reverse 5’- CTCATCTGTCCAGCTTGGGG-3’, *Dcx* forward 5’-AGGTAACGACCAAGACGCAAA-3’ and *Dcx* reverse 5’-TACCTTGTGCTTCCGCAGAC-3’. Primers targeting *Dcx* mRNA were designed to detect multiple transcript isoforms expressed from the *Dcx* gene. Primer efficiency was assayed using a standard curve (log_10_ dilution series (1:10) over a range of six concentrations (starting concentration 500-1000ng/μL). The Pfaffl method (Pfaffl, 2001) was used to determine relative expression of mRNAs with normalisation to an endogenous reference gene (ACTB) with additional within sample normalisation to ROX (a passive reference dye).

### 5.14 Statistical Analysis

GraphPad Prism (8.2.0) was used for all data analysis and to produce graphical representations. To determine the appropriate statistical test procedures for sets of observations, data were tested for parametric assumptions (i.e. Gaussian/normal distribution and sample variance equality). A Shapiro-Wilk test was used to detect normal or non-normal distribution within data sets and an F-test used to calculate whether test sample variance was significantly different from variance in the comparative control. For unpaired observations where sample variance was equal and distribution shown to be normal, a one-way ANOVA or unpaired two-tailed t-test was used to test for group differences. For unpaired observations where sample variance was not equal, but the distribution was normal, a Mann-Whitney U test was used to test group differences. For non-parametric testing of significant deviation from 0 in unpaired observations (where sample variance was not equal and the distribution not normal), a one-sample Wilcoxon signed-rank test was used to determine whether a group of observations was significantly different from 0 e.g. significant object preference in the OPS task (hypothetical mean = 0, *p<*0.05, median value of the test sample is statistically significantly different to 0). For paired data, a paired t-test or one-way ANOVA was used. For repeated measures, a repeated-measures ANOVA was used. *Post hoc* correction for multiple testing was carried out using a Bonferroni correction where indicated. Statistical significance is indicated by display of individual p-values except where denoted as *p≤0.05, **p≤0.01 and ***p≤0.001. Similarly, two-way repeated-measures Analysis of Variance (ANOVA) was used to determine factors that contribute significantly to differences observed between trials, animals and housing condition in the OPS task. For *in silico* functional genomic analysis of *Neurod1* targets, *p-*values were adjusted using the Benjamini-Hochberg *post-hoc* test to control for False Discovery Rate (FDR) < 5% (*q* < 0.05).

## 6 Results

### 6.1 Expression of miR-30e-5p is specifically increased in the ventral hippocampus after 10 weeks of social isolation during adulthood

To determine the effects of social isolation on mmu-miR-30e-5p expression, the ventral (vHIPP) and dorsal (dHIPP) hippocampus were bilaterally dissected from fresh whole hippocampi (Figure 1A and 1B) from the brains of test-naïve male BALB/c mice after 10 weeks of living either in social isolation (SI) by individual housing (n=10) or paired housing (PH) (n=8). The *Pde11a4* gene encodes a protein enriched in the vHIPP relative to dHIPP (Kelly et al., 2010), therefore to validate the dissection of distinct vHIPP from dHIPP in this study *Pde11a4* mRNA expression enrichment was used as an endogenous control. Expression of *Pde11a4* mRNA was significantly increased in the vHIPP relative to the dHIPP in a combined analysis of expression in all mice (fold-change vHIPP relative to dHIPP: [1.401, 7.785], mean fold-change = 3.964, two-tailed Wilcoxon matched-pairs signed rank test, *p<*0.0001, n=18 combined SI and PH groups, vHIPP versus dHIPP paired per animal) (Figure 1C). Comparing *Pde11a4* mRNA expression between housing groups revealed no significant difference in expression in vHIPP or dHIPP (vHIPP in SI versus vHIPP in PH or dHIPP in SI versus vHIPP in PH).

**Figure 1.**
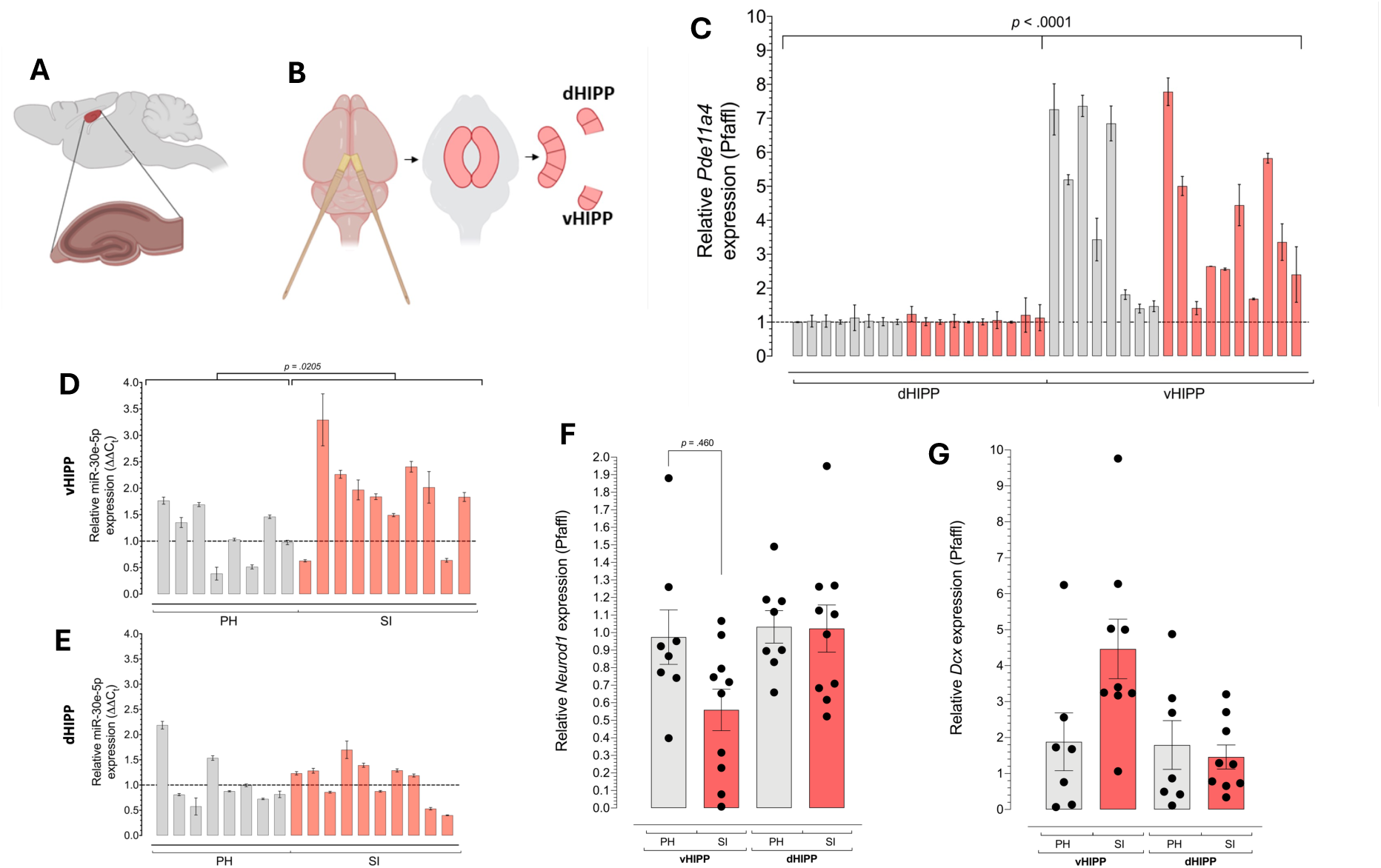
Expression of miR-30e-5p is elevated in the ventral hippocampus relative to the dorsal hippocampus in male BALB/c mice. A) Schematic of the sagittal view of a mouse brain indicating the position of the hippocampal formation within the medial temporal lobe. (B) Schematic of the manual dissection protocol used to isolate tissue from the ventral (vHIPP) and dorsal (dHIPP) hippocampal poles. (C) Quantitative PCR analysis of *Pde11a4* mRNA normalised to *Actb* mRNA and expressed as a fold-change (Pfaffl analysis) in vHIPP relative to dHIPP revealed expression of *Pde11a4* mRNA was significantly increased in the vHIPP relative to the dHIPP in all mice (grey = PH (n = 8) and red = SI (n=10)). D) and E) Quantitative PCR analysis of mmu-miR-30e-5p expression normalised to U6 small non-coding RNA and expressed as a fold-change (ΔΔCt analysis) in pair housed (PH, n=8) relative to individually housed (SI, n=10). (D) Expression was significantly higher in SI compared with PH mice in the vHIPP. E) Expression was not different in SI compared with PH mice in the dHIPP. F) Quantitative PCR analysis of *Neurod1* mRNA expression normalised to *Actb* mRNA and expressed as a fold-change (Pfaffl analysis) in pair housed (PH, n=8) relative to individually housed (SI, n=10). Expression was significantly lower in SI compared with PH mice in the vHIPP, but expression was not different in SI compared with PH mice in the dHIPP. G) Quantitative PCR analysis of *Dcx* mRNA normalised to *Actb* mRNA and expressed as a relative fold-change (Pfaffl analysis) revealed a significant increase in *Dcx* mRNA expression in SI versus PH in the vHIPP (p=0.0457) but no change in expression in the dHIPP (grey = PH (n = 7) and red = SI (n=9)). Unpaired 2-tailed *t-*test used to determine significance, threshold *p* < 0.05. Data graphed are mean ± SEM.

Next, mmu-miR-30e-5p expression was assayed and comparison between housing groups revealed expression was significantly increased in the vHIPP of SI mice relative to PH mice (SI median relative fold-change = 1.905 and mean rank = 12.10, PH median relative fold-change = 1.191 and mean rank = 6.250, Mann-Whitney *U* = 14*, p =* .0205) (Figure 1D). No significant difference in miR-30e-5p expression was observed in the dHIPP of SI relative to PH mice (SI median relative fold-change = 1.211 and mean rank = 10.00, PH median relative fold-change = 0.8461 and mean rank = 8.875, Mann-Whitney *U* = 35*, p =* .6965) (Figure 1E).

The open-access database TarBase (v8, RRID: SCR_010841, (Karagkouni et al., 2018)) was used to identify experimentally validated mmu-miR-30e-5p/mRNA interactions yielding a list of 578 target genes from one high-throughput sequencing of RNAs isolated by crosslinking immunoprecipitation (HITS-CLIP) experiment in developing mouse brain (Chi et al., 2009). One target, *Neurod1* which maps to chromosome 2 on mouse and 2q32 in humans (Tamimi et al., 1996), was identified as a gene target of interest since it was associated with the GOBP database term “neurogenesis” (GO:0022008, AmiGO, RRID: SCR_002143 and GeneCards, RRID: SCR_002773) and is reported in the literature as a regulator of neurogenesis and brain development (Boutin et al., 2010; Gao et al., 2009; Liu et al., 2020; Schwab et al., 2000; Yu et al., 2014). *Neurod1* encodes a transcription factor targeting genes enriched in “neurogenesis” (GO:0022008) (TRRUST) (Supplementary Data 1) and other neurogenesis-related GOBP terms (Supplementary Data 2). Additionally, it targets multiple signalling pathways (KEGG pathways, RRID: SCR_012773, Supplementary Data 3).

Therefore, we determined expression of *Neurod1* mRNA in vHIPP and dHIPP. *Neurod1* mRNA was significantly decreased in the vHIPP of SI mice (n=10) relative to PH mice (n=8) (unpaired 2-tailed *t-*test, *p* = 0.0460, n=8 PH, n = 10 SI) (Figure 1F), but no change was observed in the dHIPP (SI versus PH) (unpaired 2-tailed *t-*test, *p* = 0.9559, n=8 PH, n = 10 SI) (1E). Expression of *Neurod1* mRNA was not significantly different in the vHIPP relative to dHIPP in a combined group of all mice (unpaired 2-tailed *t-*test, *p* > 0.05, n=18 vHIPP versus n=18 dHIPP). Expression of the neuroblast marker *Dcx* mRNA was assayed as a molecular indicator of neurogenesis. This was done in 7 SI and 9 PH test-naïve mice due to insufficient RNA sample from 2 mice (1 from each housing group). In the vHIPP, expression of *Dcx* mRNA was significantly increased in SI mice compared with PH mice (unpaired 2-tailed *t-*test, *p* = 0.0457, n=7 PH, n = 9 SI) but not dHIPP (unpaired 2-tailed *t*-test, *p* = 0.6440, n=7 PH, n = 9 SI) (Figure 1G).

### 6.2 Social isolation during adulthood was not associated with significant change to affective-like behaviours

Anxiety-like behaviour was measured in the open-field test. No difference was observed in overall locomotion between individually housed (SI, n=10) and pair housed (PH, n=10) mice as measured by comparing velocity of SI and PH mice in the whole arena (SI mean velocity in whole arena (m s^-1^) = 3.421, SD = 0.959, PH mean velocity in whole arena (m s^-1^) = 3.636, SD = 1.172, unpaired 2-tailed *t-test, F* [9,9] = 1.493*, p =* .6588) and in the outer zone only (SI mean velocity in outer zone (m s^-1^) = 3.25, SD = 0.8612, PH mean velocity in outer zone (m s^-1^) = 3.564, SD = 1.246, unpaired 2-tailed *t-test, F*[9,9] = 2.092*, p =* .5203). Total distance travelled in the arena was comparable between SI and PH mice (SI mean distance (cm) = 2032.00, SD = 560.1, PH mean distance (cm) = 2155.00, SD = 679.90, unpaired 2-tailed *t-test, F* [9,9] = 1.473*, p =* .6638) and in the outer zone only (SI mean distance (cm) = 1734.00, SD = 443.60, PH mean distance (cm) = 1378.00, SD = 382.70, unpaired 2-tailed *t-test, F* [9,9] = 1.343*, p =* .0703).

Qualitative comparison of movement traces of individually housed mice (n=10) to pair-housed mice (n=10) initially indicated SI mice avoid the central zone whereas PH mice explore the whole arena (Figure 2A). However, no significant difference was observed in distance travelled in the centre zone (SI median distance (cm) = 55.43 and mean rank = 8.80, PH median distance (cm) = 323.8 and mean rank = 12.20, mean rank = 11.20, Mann-Whitney *U* = 33*, p =* .2168) (Figure 2B), neither in the number of entries into the centre zone (SI median entries = 4.50 and mean rank = 8.80, PH median entries = 27.00, mean rank = 12.20, Mann-Whitney *U* = 33*, p =* .2085) (Figure 2C) nor in the duration spent in the centre zone (SI median time (s) = 9.88 and mean rank = 9.10, PH median time (s) = 40.20 and mean rank = 11.90, mean rank = 11.20, Mann-Whitney *U* = 36*, p =* .3140) (Figure 2D). No difference was observed in the latency to enter in the centre zone between SI and PH mice (SI median latency (s) = 10.76 and mean rank = 8.90, PH median latency (s) = 29.80, mean rank = 12.10, Mann-Whitney *U* = 34*, p =* .2390) (Figure 2E) in SI (n=10) compared with PH (n=10) mice. In the latency to enter the centre zone, 2 PH mice and 1 SI mouse did not move from the outer zone of the arena at any point in the 10-minute trial and so were identified as statistical outliers (ROUT test, Q=0.1%). Another SI mouse was identified as a statistical outlier in the number of entries, distance travelled and duration in centre zone (ROUT test, Q=0.1%). Outliers were retained in the data sets and indicated in the graphs presented (outlier data points = × on the graphs in Figures 2B, 2C, 2D and 2E).

**Figure 2.**
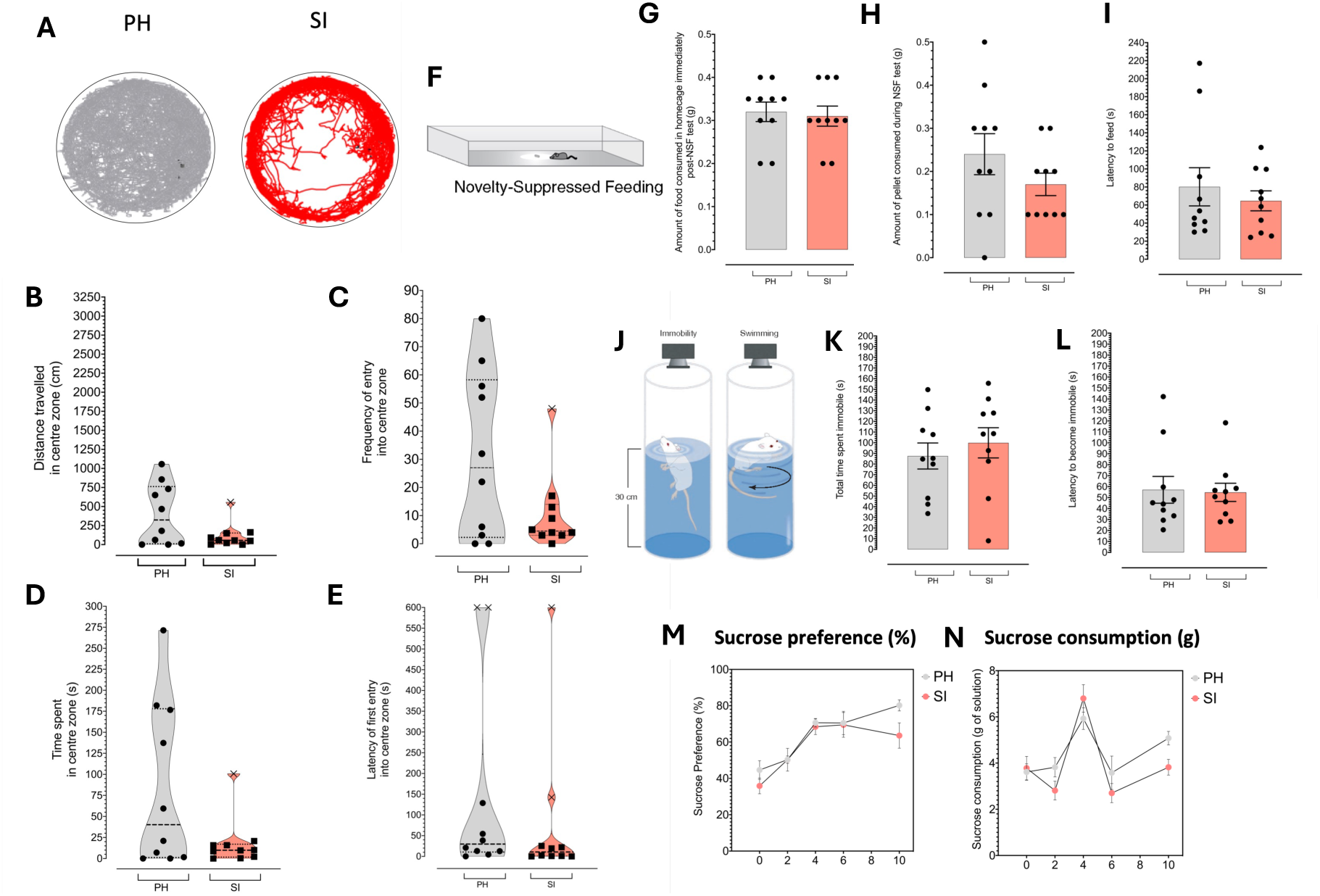
Measure of open field avoidance in male BALB/c mice housing in social isolation. (A) Traces overlaid on arena schematic for each mouse (PH, left panel, n = 10 and SI, right panel, n=10). (B) Distance travelled, (C) frequency of entry, (D) time spent (duration) and (E) latency to enter in centre zone was not different between PH and SI mice. Each animal is represented by individual data points graphed (data points for statistical outliers in each measure are denoted using symbol ×). Violin plots graph median (central dashed line), interquartile range (dotted lines above and below dashed line) and range (solid line at top and bottom of each plot) for each dataset with graphical display of dataset distribution. F) Schematic of NSF test arena into which the mouse is placed after 24h food restriction (no access to food but water ad libitum). (H) Amount of food consumed in the 5 mins immediately post-test upon return to home cage, (I) latency to feed was not different in mice housed individually (SI, n=10) compared with pair-housed mice (PH, n =10). (J) Schematic of the apparatus used for the Porsolt swim test in which the mouse is placed in a clear Perspex cylinder (25 cm diameter) and the time spent immobile (left panel), the time taken for the mouse to first become immobile versus swimming (right panel) scored post-acquisition from a DVD recording. No difference was observed in (K) total immobility time or (L) latency to become immobile in mice housed individually (SI, n = 10) compared with pair-housed mice. Graphical representations of (M) sucrose preference (%) and (N) absolute consumption (g of solution) which was not statistically different between individually and pair-housed mice among the tested mice (SI, n = 10 and PH, n = 10, sucrose preference/consumption was measured in week 0, 2, 4, 6 and 10) or test-naïve mice (SI, n = 10 and PH, n = 10, sucrose preference/consumption was measured in week 0, 2, 4 and 6). Data graphed are mean ± SEM; each animal is represented by a data point on each graph (except M and N where datapoint = time)

Depression-like behaviour was tested using a combination of typical tests. The novelty suppressed feeding (NSF) test was used to measure of novelty induced anxiety/depression-like hyponeophagia, a test widely conducted to determine anxiety-like behaviour in rodents and is established as sensitive to antidepressant treatment (Samuels and Hen, 2011). The NSF test was conducted after 10 weeks of social isolation and no significant difference in NSF behaviour was observed in individually housed mice relative to pair-housed mice for post-test homecage refeeding, amount of pellet consumed or latency to feed (Figure 2G, 2H and 2I respectively). The Porsolt Swim Test is a widely used measure of behavioural despair (Figure 2J). In the test, conducted in week 10, no difference was observed in immobility time (Figure 2K) or latency to become immobile between groups (Figure 2L).

Grooming behaviour was assessed by cumulative coat state measure taken weekly simultaneous with body weight and food/water consumption measures throughout testing for all animals, and no difference in coat state was observed in SI versus PH mice at any stage. Food or water consumption was not significantly different between groups. No differences were observed in body weight (Supplementary Figure 4A). Additionally, latency to groom and time spent grooming after coat soiling was similar between groups (Supplementary Figure 4B). Olfaction was intact in all animals as determined by the cookie test (Supplementary Figure 2D) and outcomes from the social dominance (Supplementary Figure 2E) test did not indicate significant dominance interaction between cagemates in PH mice.

The sucrose preference/consumption test is a widely used as a measure of anhedonia conducted at the start of the experiment (week 0) and after 2, 4, 6 and 10 weeks of individual or paired housing. No significant difference was observed in sucrose preference (Figure 2M) at Week 0 (SI median preference = 40.31%, PH median preference = 38.15, Mann-Whitney *U* = 180, *p* = .5968, SI n = 20, PH n=18) or after 2 weeks (SI median preference = 55.21%, PH median preference = 48.40, Mann-Whitney *U* = 136, *p* = .0845, SI n = 20, PH n=18), 4 weeks (SI median preference = 68.65%, PH median preference = 69.48, Mann-Whitney *U* = 156, *p* = .4917, SI n = 20, PH n=20), 6 weeks (SI median preference = 74.80%, PH median preference = 83.15, Mann-Whitney *U* = 172, *p* = .4567, SI n = 20, PH n=20) or 10 weeks (SI median preference = 63.40%, PH median preference = 83.60, Mann-Whitney *U* = 26, *p* = .0715, SI n = 10, PH n=10) of individual housing relative to paired housing. Absolute sucrose consumption (Figure 2N) was similar in SI versus PH mice in week 0, 2, 4 and 6, but in week 10 sucrose consumption was significantly lower in SI mice compared with PH mice (SI median sucrose consumption (g) = 3.90, PH median sucrose consumption (g) = 4.80, Mann-Whitney *U* = 16, *p* = .0083, SI n = 10, PH n=10) although this difference was not significant after controlling for false-discovery rate (Benjamini-Hochberg method, *q* = 0.0597).

### 6.3 Social isolation during adulthood has no effect on basal circulating CORT levels but attenuates acute stress induced CORT increase

To determine if individually housed male BALB/c mice presented with elevated levels of endogenous glucocorticoids, circulating basal levels of the main murine glucocorticoid, CORT, was quantified from plasma samples collected in the early evening (all animals sampled between 1700 and 1800 hours after 10 weeks of pair or individual housing immediately prior to sacrifice). Comparing a combined group of tested and test naïve individually housed mice with a combined group of pair housed mice, no difference was found in plasma levels of CORT (SI median [CORT]ng/μL = 16.12, PH median [CORT]ng/μL = 29.77, Mann-Whitney *U* = 128.5, *p* = .1353, SI n = 20, PH n=18, Figure 3A). Comparing tested individually house mice with tested pair housed mice, no difference was found in plasma levels of CORT (SI median [CORT]ng/μL = 15.93, PH median [CORT]ng/μL = 21.89, Mann-Whitney *U* = 33.5, *p* = .2176, SI n = 10, PH n=10). Comparing test naïve individually housed mice with test naïve pair housed mice, no difference was found in plasma levels of CORT (SI median [CORT]ng/μL = 20.08, PH median [CORT]ng/μL = 44.87, Mann-Whitney *U* = 28, *p* = .3154, SI n = 10, PH n=8). No difference was observed in plasma CORT concentration in all tested versus test naïve mice (combined SI and PH, tested median = 18.46, test naïve median = 29.90, Mann-Whitney *U* = 131.5, *p* = .1601, SI n =20, PH n = 18), neither in SI tested mice versus SI test naïve mice (SI tested median = 15.93, SI test naïve median = 20.08, Mann-Whitney *U* = 36, *p* = .3150, SI tested n =10, SI test naïve n = 10) nor PH tested mice versus PH test naïve mice (PH tested median = 21.89, PH test naïve median = 44.87, Mann-Whitney *U* = 27, *p* = .2743, PH tested n =10, PH test naïve n = 8). Plasma CORT concentration was quantified from samples collected 24h prior to the Porsolt Swim test in the 10^th^ week as a measure of CORT levels prior to an acute stressor. No difference in plasma CORT concentration was observed when comparing individually housed with pair housed mice (SI median = 20.31, PH median = 46.24, Mann-Whitney *U* = 26, *p* = .2278, SI n =10, PH n = 8). Plasma CORT concentration was quantified from samples collected 30 mins post-Porsolt Swim test in the 10^th^ week as a measure of CORT levels immediately after an acute stressor. Comparison of plasma CORT concentration 24h prior to and 30 mins after the Porsolt Swim test for all mice revealed a significant rise in circulating CORT 30 mins post-test (combined SI and PH median pre-test [CORT]ng/μL = 20.31, median post-test [CORT]ng/μL = 72.97, Wilcoxon matched-pairs signed rank test, *p* < 0.0001). Within housing group comparison revealed plasma CORT concentration was significantly elevated in pre-and post-test measures in individually housed mice (SI median pre-test [CORT]ng/μL = 20.31, SI median post-test [CORT]ng/μL = 72.97, Wilcoxon matched-pairs signed rank test, *p* = .0137) and in pair housed mice (PH median pre-test [CORT]ng/μL = 60.51, PH median post-test [CORT]ng/μL = 165.0, Wilcoxon matched-pairs signed rank test, *p* = .0078, Figure 3B). However, the post-stress increase in plasma CORT concentration in individual housed mice was significantly lower relative to pair housed mice (SI median = 20.31, PH median = 46.24, Mann-Whitney *U* = 26, *p* = .2278, SI n =10, PH n = 8, Figure 3B).

**Figure 3.**
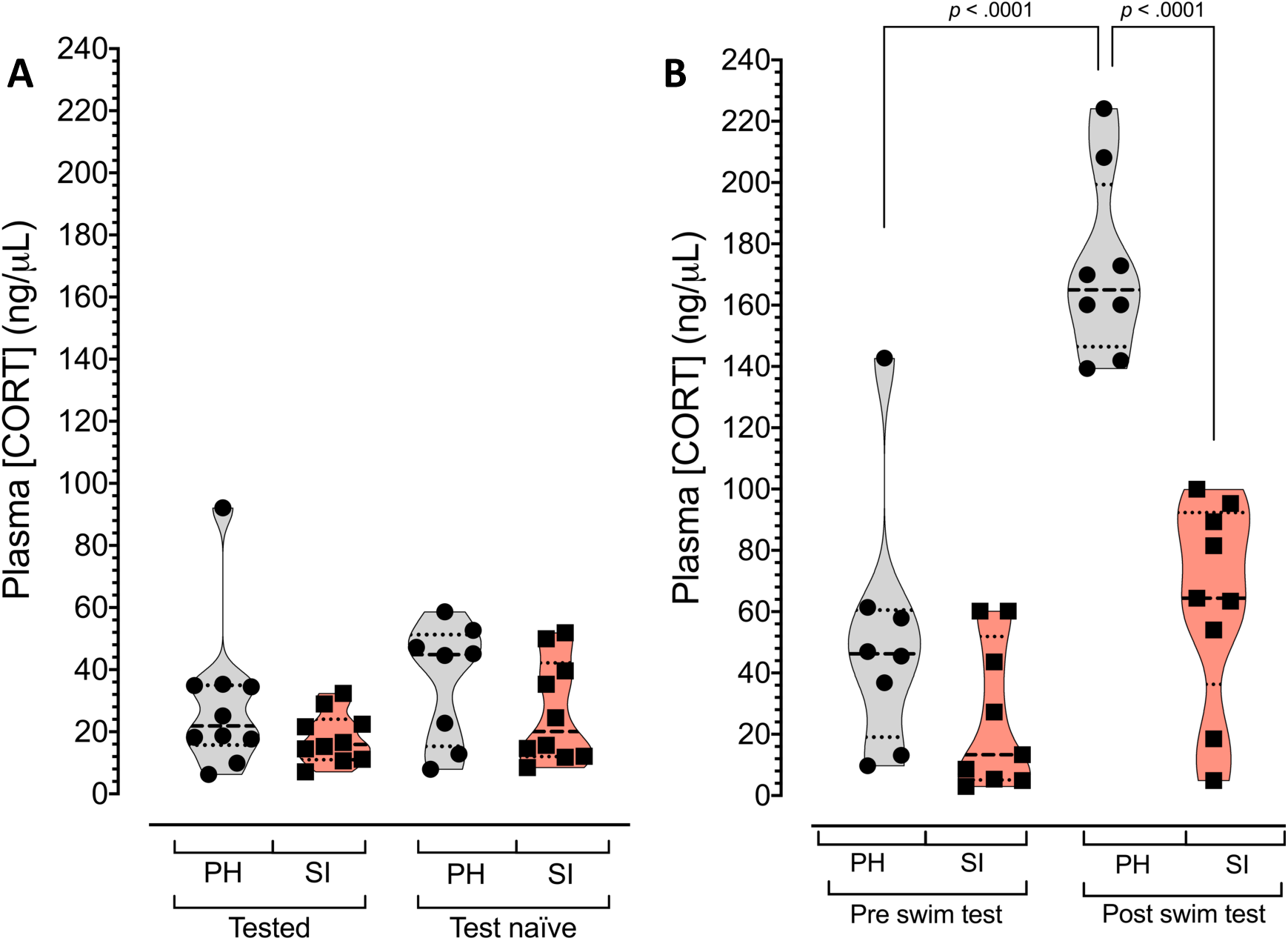
Measure of circulating corticosterone concentration from plasma samples collected from male BALB/c mice after 10 weeks of social isolation by individual housing. (A) Basal CORT concentration was determined from plasma samples collected in the evening over a narrow time window and quantified by ELISA (no significant difference was observed). (B) CORT concentration was also determined 24h pre Porsolt swim test, and 30 mins post swim test to quantify plasma CORT levels prior to and after an acutely stressful event. CORT was significantly increased in PH mice post swim test compared with pre swim test, whereas no difference was found in SI mice in pre versus post swim test plasma CORT concentration. Between housing groups, CORT levels were comparable in PH and SI mice pre swim test, but CORT levels in SI mice was significant reduced relative to PH mice post swim test. Violin plots graph median (central dashed line), interquartile range (dotted lines above and below dashed line) and range (solid line at top and bottom of each plot) for each dataset with graphical display of dataset distribution.

### 6.4 Object exploration behaviour in object pattern separation task is altered after 10 weeks of social isolation adult male mice cognition

In order to test adult hippocampal neurogenesis-dependent cognition, we used an object pattern separation (OPS) task (adapted from (van Hagen et al., 2015)), which is a variant of the object recognition tasks commonly conducted in rodents to test contextual cognition. The OPS test arena contained two identical objects for exploration (Figure 4A). Over the course of 4 trials (T1-4) one object was displaced to novel positions within the arena whilst the other remained static in T1-4 (Figure 4B).

**Figure 4.**
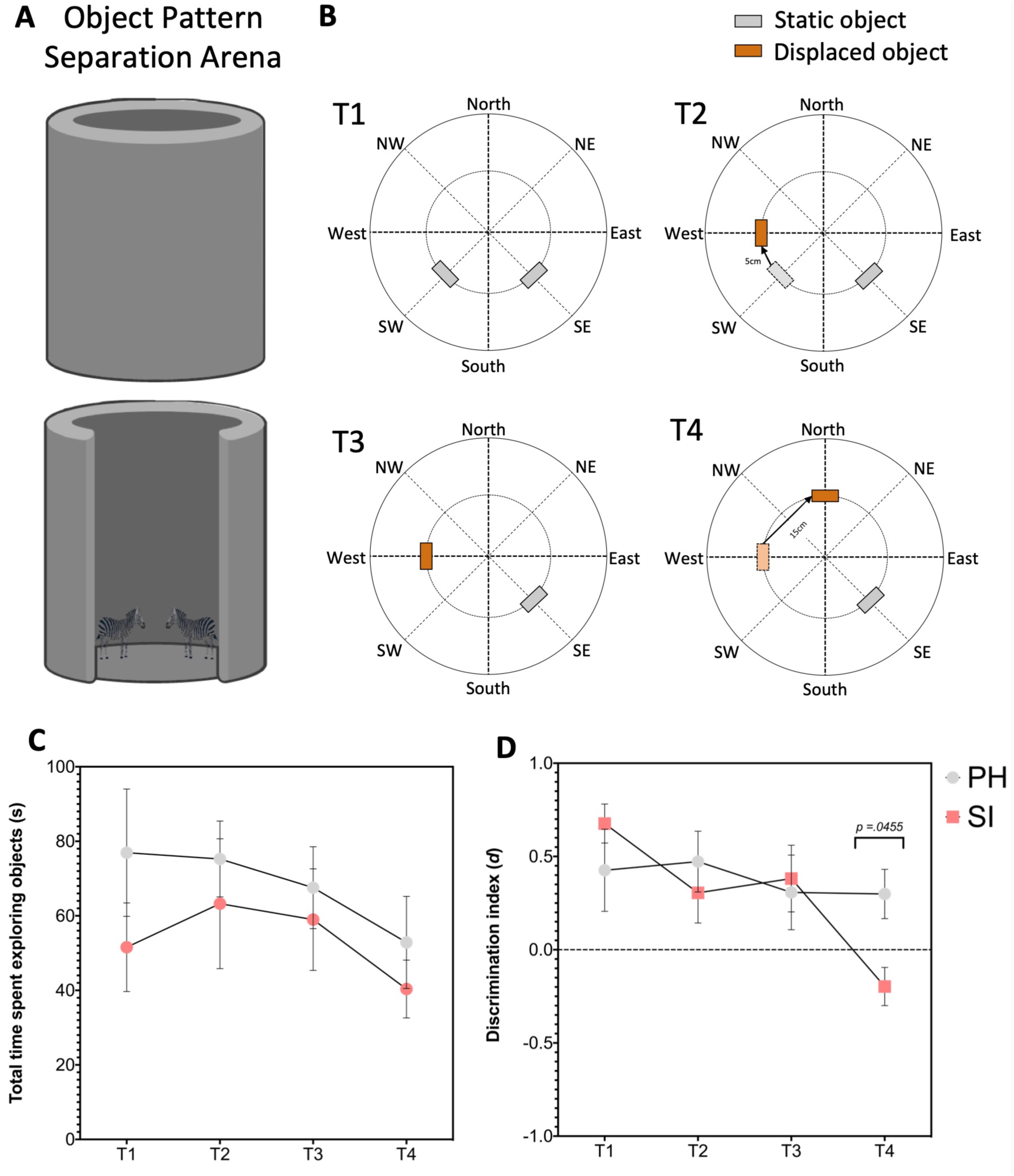
Measure of pattern separation-like cognition by object pattern separation test in male BALB/c mice after social isolation by individual housing. (A) Schematic of the test arena consisting of the same container as the open field test (40cm diameter, walls 40cm high) containing objects for exploration by each animal. (B) Schematics outlining the object positions in T1-4. (C) Time spent exploring the objects was calculated as the sum of object exploration for each object per trial, no difference was observed between housing groups. (D) The discrimination index (*d*) was calculated to express relative object exploration as a number between -1 and 1 and no difference was found between housing groups in T1-3 but a significant decrease in relative object exploration as the object was displaced 15cm in T4 was observed in SI (n =7) versus PH mice (n = 8). Mice were excluded if they showed a lack of exploratory behaviour as determined by either no interaction with one or both objects in any trial or a lack of open field and object exploration during habituation.

Total time spent exploring (sum of object exploration for both objects was calculated for each trial) by mice in SI and PH was similar in each trial (Figure 4C) and there were no significant differences in overall exploratory behaviour as a result of housing (two-way *ANOVA, F* [1,17] = 3.276, % of total variation = 8.058, *p =* .0880), trial or (two-way *ANOVA, F* [3,51] = 2.509, % of total variation = 6.350, *p =* .691) or housing × trial combined (two-way *ANOVA, F* [3,51] = 0.3138, % of total variation = 0.7941, *p =* .8153). This suggests housing in SI compared with PH in male BALB/c mice has no impact on general object exploratory behaviour, although individual exploratory phenotype varies significantly between animals.

A discrimination index (*d*) was used to compare relative object preference across trials between housing groups. There was a significant effect of trial × housing condition combined (two-way *ANOVA, F* [3,39] = 2.902, % of total variation = 8.481, *p* = .0469), trial (two-way *ANOVA, F* [2.115, 27.49] = 4.829, % of total variation = 14.11, *p* =.0146). There was no significant effect of housing condition alone (two-way *ANOVA, F* [1,13] = 0.2525, % of total variation = 0.7748, *p* = .6237). Group-wise comparison revealed no significant difference between *d* in SI compared with PH mice in T1-T3 (Bonferroni *post hoc* correction for multiple testing, n = 8 PH and n = 7 SI, **TI:** PH mean = 0.4260 and SI mean = 0.6768, *p* >.9999, **T2:** PH mean = 0.4727 and SI mean = 0.3050, *p* >.9999, **T3:** PH mean = 0.3074 and SI mean = 0.3816, *p* >.9999) (Figure 4D). In T4, there was a significant decrease in *d* in SI relative to PH mice (Bonferroni *post hoc* correction for multiple testing, n = 8 PH and n = 7 SI, **T4:** PH mean = 0.2987 and SI mean = -0.1973, *p* = .446) (Figure 4D).

Because a significant between-trial effect was observed, comparison was made across trials within each housing group. No difference was observed in *d* for any between-trial comparison in PH mice (Bonferroni *post hoc* correction for multiple testing, n = 8 PH, **T1 v T2:** *p* > .9999, **T1 v T3:** *p* > .9999, **T1 v T4:** *p* > .9999, **T2 v T3:** *p* = .7133, **T2 v T4:** *p* > .9999, **T3 v T4:** *p* > .9999). In SI mice however, a significant decrease in relative exploration of the displaced object was observed in T1 versus T4 (Bonferroni *post hoc* correction for multiple testing, n = 8 PH, **T1 v T2:** *p* = .2054, **T1 v T3:** *p* = .7704, **T1 v T4:** *p =* .0045, **T2 v T3:** *p* > .9999, **T2 v T4:** *p* = .5040, **T3 v T4:** *p* = .0657).

## 7 Discussion

Social isolation was associated with significant changes to gene expression and modest changes to object pattern separation and stress. Increased expression of the miRNA mmu-miR-30e-5p was accompanied by a decreased in *Neurod1* mRNA, a neurogenesis-related transcription factor, in the ventral hippocampus in individually housed mice compared with those housed in pairs. Increased anxiety and/or affective-like phenotypes were expected in individually housed mice relative to pair housed mice, but there were no significant changes observed in the open field test for anxiety-like behaviour, NSF and sucrose preference/consumption for anhedonia and Porsolt swim test for behavioural despair. In a test of pattern separation-like cognition, socially isolated mice appeared to lose preference for exploration of an object in a novel location despite initially showing similar exploratory behaviour to pair housed mice.

The social isolation paradigm used, where mice are housed individually rather than in pairs, is a unique approach since previous studies compared socially isolated mice to groups of mice housed with as many as 15 cagemates (e.g. (Mudra Rakshasa and Tong, 2020)). Group housed male mice are likely to be stressed given the establishment of social hierarchies through aggressive behaviour seen between male mice (Brown, 1953; Weber et al., 2022). Moreover, a majority of previous reports investigating social isolation by individual housing have been done earlier in development (post-weaning), the equivalent of early adolescence) (e.g. (McGrath and Briand, 2022; Myers et al., 2024). In studies which investigate adult rodents, social isolation in rats or transgenic animals, rather than inbred mice, are more widely studied (Kim et al., 2025; Powell and Swerdlow, 2023; Toyoshima and Yamada, 2023). In addition, tissue samples were carefully and separately dissected from the dorsal and ventral hippocampus using a previously established protocol (Kelly et al., 2010), given that the dorsal and ventral hippocampus are functionally and neuroanatomically distinct substructures (Fanselow and Dong, 2010; Henke, 1990; Moser and Moser, 1998). No other study, to our best knowledge, report on microRNA expression changes in the sub-regions of the hippocampus of socially isolated adult mice making the present study the first to use a social isolation model to examine cognition, anxiety/depression-like behaviours and microRNA-mRNA expression in the dorsal and ventral hippocampus of adult mice.

Expression of miR-30e is associated with psychiatric disorders. For example, expression of miR-30e is altered in plasma, peripheral mononuclear blood cells and post-mortem tissue from individuals with psychiatric disorders (Gorinski et al., 2019; Kohen et al., 2014; Malekpour et al., 2026; Perkins et al., 2007; Sun et al., 2015). Genetic studies had previously demonstrated a variant in the precursor of hsa-miR-30e-5p alters the expression of the mature sequence and is linked with MDD and cognitive dysfunction (Xu et al., 2010b). Overexpression of miR-30e-5p in the hippocampus of adult rats has been shown to impair cognition (Xu et al., 2015) due to aberrant brain connectivity using resting state fMRI, with antidepressant drugs partially reversing cognitive impairment in this model (Xu et al., 2016). Therefore, we proposed that altered expression of mmu-miR-30e-5p would be associated with altered cognitive function in a mouse model of social isolation in adulthood.

We demonstrated an increase in miR-30e-5p expression in the ventral hippocampus of socially isolated mice in comparison to paired mice. However, in a previous study, miR-30e expression was significantly decreased along with multiple other members of the miR-30 family in the DG of adult mice after chronic defeat stress (Khandelwal et al., 2019). The contrasting findings could be due to the use of the different stress paradigms which includes both psychological and physical stressors. Repeated subordination by exposure to a dominant, aggressor mouse could itself result in depressive-like phenotype (Golden et al., 2011) via an alternative mechanism involving miR-30e. Additionally, Khandelwal et al., (2019) isolated the DG specifically via needle-punch without separating samples at the co-ordinates relative to Bregma along the dorso-ventral axis and did not specify whether it was miR-30e-5p or miR-30e-3p (alternative mature forms of the same pre-miRNA but with different target gene sets) was assayed. In another study, a related miRNA from the miR-30 family, miR-30a-5p, was transiently down-regulated at postnatal day 21, followed by a significant upregulation at postnatal day 40, which was sustained until at postnatal day 62 in prenatal stress (Cattaneo et al., 2020) therefore age at time of stress and experimental procedures could also play role. Thus, factors such as age, tissue isolation method, the form of the mature miRNA being measured, and the type of stress could also impact the expression levels of miRNA-30 family including mmu-miR-30e-5p.

To validate mmu-miR-30e-5p regulatory targets in the current study, the expression of *Nrp1* and *Neurod1* mRNA were assayed in the samples extracted from both the dorsal and ventral hippocampus and the differences between individually and pair-housed mice were compared. No difference in *Nrp1* mRNA expression was observed, indicating molecular regulation in the hippocampus of the current study does not involve significant changes in expression of *Nrp1* as it does in the DG of mice after chronic social defeat stress in Khandelwal et al (2019). Additionally, *Neurod1* stood out as a candidate since it has an established role in neurogenesis in mouse and non-human primates (Fedele et al., 2011; Gao et al., 2009; Tutukova et al., 2021). The results of a HITS-CLIP experiment in developing mouse brain, made available for *in silico* data mining (Karagkouni et al., 2018), showed *Neurod1* mRNA is a target of mmu-miR-30e-5p (Chi et al., 2009). In this current study, it was discovered that expression of *Neurod1* mRNA was significantly decreased in the ventral, but not dorsal, hippocampus. Functional genomic analysis suggests the downstream genomic targets of *Neurod1* are enriched in the GOBP terms associated with neurogenesis (“neuron migration” GO:0001764, “axon guidance” GO:0007411, “brain development” GO:0007420 and “nervous system development” GO:0007399) and other relevant pathways identified in the KEGG database.

The possibility that mmu-miR-30e-5p/Neurod1 mRNA regulatory interaction involved direct interaction should be further investigated as this interaction was predicted *in silico* on the basis of existing data(Chi et al., 2009; Karagkouni et al., 2018) in this study, rather than tested experimentally. The canonical function of miRNAs is to act as guides that recruit RNA-induced silencing complex (RISC) to specific target mRNAs to repress protein translation. Argonaute 2 (Ago2) is the miRNA interacting component of the RISC, therefore, Ago2-RNA immunoprecipitation was used to isolate the miRNA/mRNA interacting fraction in this study (data not shown). However, the total RNA isolated from a small portion of dorsal and ventral hippocampal tissue using immunoprecipitation was insufficient for gene expression analysis despite the sensitivity of qPCR. In future, such analysis would yield important insights into the endogenous miRNA/mRNA interactome *in vivo* but it would likely require pooled samples from multiple replicates to generate sufficient RNA yield. The functional relevance of the regulatory interaction between mmu-miR-30e-5p and *Neurod1* mRNA would require confirmation with a reverse genetic study combining knockdown and overexpression of both mmu-miR-30e-5p and *Neurod1* in the adult dorsal and ventral hippocampus. The functional genomic analysis hints that targets of Neurod1 transcription factor, encoded by *Neurod1* mRNA, were enriched in biological processes associated with hippocampal neurogenesis, however these targets should be confirmed using ChIP-sequencing in future studies.

Hippocampal neurogenesis was not examined in this currentstudy. Thus, histological techniques coupled with the behavioural and molecular experiments could help clarify the connection between the inverse relationship between mmu-miR-30e-5p and *Neurod1* and neurogenesis markers (for example a BrdU-incorporation assay). To fully understand whether mmu-miR-30e-5p regulation in the ventral hippocampus is linked with DG granule neurons specifically, it would be compelling to carry out single cell expression analysis in human and mouse DG granule neurons.

Whether the altered gene expression reported here occurs along with robust or significant changes to behaviour and/or cognition in social isolation is unclear. In terms of anxiety-like behaviour, no significant difference was observed in any quantitative measure of open-field avoidance or locomotion in the open field test between the two groups of differently housed mice. A common behavioural trait reported in models of social isolation is an increase in thigmotaxis as determined by open field testing (Burke et al., 2025; Mumtaz et al., 2018). However, we observed no significant difference in the open field avoidance test between housing groups after 6 weeks of isolation. Statistical outliers skewed quantitative measures, but no outliers were removed from the datasets post hoc. Studies also describe generalised hyperactivity as a feature of social isolation (Einon and Morgan, 1978; Sullens et al., 2021). Although, general hyperactivity or increased locomotion in the open field test was not observed in this study. Furthermore, the NSF test, conducted at 10 weeks, found no differences between groups in anxiety/depression-like novelty induced hyponeophagia. Anxiety-like behaviour in rodents is frequently associated with depression-like behaviours such as anhedonia and behavioural despair and combined testing of both anxiety and depression-like behaviours is typical in investigating rodent models of MDD (Petković and Chaudhury, 2022). Therefore, we did not confirm that social isolation by individual housing in adulthood is associated with significant change in anxiety-like behaviour in male BALB/c mice. Perhaps extending the social isolation period would induce a phenotype that is more robustly distinguishable. Another key difference between the current study and previous studies is the age at which mice were socially isolated. Social isolation may have a more significant impact on anxiety and depression-like behaviours when mice are isolated earlier in development. Extending the length of isolation may be needed to induce a response in resilient mice that represent as outlier non-responders in the current study. Increased sample sizes for greater experimental power would be needed to confirm given the variation and presence of outliers in the data.

Such variation in behavioural measures in laboratory rodents may occur due to individual differences in response to a change in social environment despite the highly similar or identical genetic background in inbred mouse lines (Beery and Kaufer, 2015; Jakovcevski et al., 2008). Since the mice used in this study were purchased as adult animals, we could not monitor or control for early life experiences in their breeding facility which can also contribute to phenotypic variability in behavioural response to our social isolation model.

A previous report from our research group showed male BALB/c mice housed individually showed significant changes in measures the Porsolt swim test and sucrose preference/consumption test (Du Preez et al., 2021). In contrast, we observed no significant differences in either test in the current study. Such results could be due to test experience given the this is the first time the OPS test was included within the battery of behavioural testing in our research group.

Sample size may also be an issue given the small effect sizes seen in the sucrose preference and NSF tests. Indeed, in the previous publication from our group with double the sample size as used in this study (n = 20 mice per housing group (Du Preez et al., 2020) found a significant change in stress free controls and stressed mice.

Previous studies measuring object pattern separation, report a preference for object exploration of the displaced object only in male mice that were group-housed for 2, 4 and 8 weeks (Chen et al., 2016). Similarly, whilst we observed no major differences between housing groups T1-3, a trend suggests that after a series of object displacements (5cm in T2 and 15cm in T4) pair housed mice sustained a preference for exploration of the object in a novel location. Relative exploration of the objects was similar in individually housed relative to pair-housed mice until T4 when individually housed mice showed a significant decline in exploration preference for the object in the novel location compared with pair-housed mice. The mechanism which drives this difference between housing groups in the final trial could be associated with altered neurogenesis.

Wereported an increase in *Dcx* mRNA in the vHIPP, consistent with a previous study in the same model of social isolation which reported increased density of very early, post-mitotic immature Dcx+ neurons after classification of Dcx+ neuroblasts based on morphology in the vHIPP, whilst other classes of Dcx+ neuroblasts remained similar to controls (Du Preez et al., 2020). Increased neurogenesis the vHIPP is important for improving cognition, specifically improving flexibility, and therefore has a proposed role in mood regulation (Anacker and Hen, 2017). Our gene expression assay was designed to detect multiple transcript isoforms expressed from the *Dcx* gene, and it has emerged that splice differences and isoforms are functionally distinct with proposed roles in tubulin dynamics and migration (Gleeson et al., 1999; Manka and Moores, 2020). Differences may have been driven in a specific splice variant of the mRNA. Future studies could benefit from investigating the correlation between isoforms of Dcx, neurogenesis and object pattern separation cognition outcomes within the same mice to determine the interaction between neurogenesis and cognition after social isolation during adulthood. Additionally exploring neurite outgrowth and migration of neuroblasts within the DG after social isolation would provide deeper phenotypic insights into the role of *Dcx* in response to social isolation.

Stress, measured by assay of plasma corticosterone (CORT) concentration at different time points, demonstrated no significant differences in basal circulating CORT levels, however, there was a significant increase in CORT following the Porsolt swim test (an acute stressor) in mice housed in paired only. This suggests a blunting of the typical CORT response to an acute stressor in socially isolated mice. Social isolation is considered a form of mild psychological stress, but it is also not necessarily associated with increased basal levels of circulating glucocorticoids, as exemplified in few previous studies that report no changes in basal CORT levels after social isolation (Arndt et al., 2009; Bartolomucci et al., 2003). Another study indicated decreased basal CORT in male C57BL/6J mice after a period of social isolation in adulthood (Ieraci et al., 2016), however, this could be due to circadian fluctuations in CORT levels.

In healthy humans, cortisol is highest in the morning and declines through the day (Chan and Debono, 2010). Conversely, since rodents are nocturnal, they display the opposite relationship with CORT levels being lowest in the morning and being highest in the late afternoon/early evening (James et al., 2023; Kalsbeek et al., 2003). In a model of acute isolation stress, group-housed (n = 15) mice were transferred to individual housing for 12h (Takatsu-Coleman et al., 2013) and the authors reported depression-like behaviour and increased plasma CORT levels in isolated animals relative to controls. Perhaps therefore, an initial acute stress response occurs in the immediate hours or days of social isolation, but this was not tested in the current study since it was the aim to investigate the effects of a prolonged period of social isolation in adulthood rather than acute effects. Interestingly, plasma CORT levels in the isolation group (167.5±15.34 ng/mL) and in the crowded group (136.0±8.43 ng/mL) were both significantly higher than the control group (76.7±7.24 ng/mL) but not different from each other (Takatsu-Coleman et al., 2013) suggesting overcrowding is as stressful as isolation. But in the same study, the overcrowded group of mice did not exhibit mood-related behavioural abnormalities as were seen in the isolated group despite having similar circulating CORT levels. This suggests that the impact of isolation is not due to altering basal circulating CORT levels but perhaps altering the typical CORT response to acute stress as found previously (Du Preez et al., 2020). Previous studies have found that social isolation in rodents leads to a dysregulation of the hypothalamic-pituitary-adrenal (HPA) axis, often resulting in exaggerated or blunted CORT levels depending on the age of isolation, duration, and sex of the mice (Brandt et al., 2022).

The use of male mice only is an important limitation. Although this approach reduced sex-related variability in this initial study, it limits the generalisability of the findings. Given known sex differences in affective disorder vulnerability, HPA-axis regulation, hippocampal plasticity and microRNA expression, future work should test whether the miR-30e-5p/Neurod1 changes and pattern separation deficits observed here are reproduced in females

In neurogenesis-ablated adult mice, CORT levels were significantly lower after acute physical stress but CORT levels in basal conditions were similar between neurogenesis ablated and wild-type mice (Snyder et al., 2011). These findings together suggest that social isolation in adult male BALB/c mice may not be associated with a chronic stress response (i.e. high basal CORT levels), however, the response to an acute stressor is altered in individually housed male BALB/c mice compared with pair housed mice via putative mechanisms involving adult hippocampal neurogenesis. Future studies where CORT levels are monitored over 24hrs could determine if disturbance to circadian fluctuations occurs in social isolation in mice in adulthood.

Collectively, these findings suggest that prolonged adult social isolation preferentially alters ventral hippocampal gene regulation and stress response, with modest effects on pattern separation-like cognition but limited impact on anxiety- or depression-like behaviours. The data support a model in which miRNA-transcription factor interactions, particularly involving miR-30e-5p and *Neurod1,* contribute to region-specific transcriptional adaptations. More broadly, the results align with emerging evidence that microRNAs, including miR-30 family members as well as miR-124-3p and miR-137, may function as non-redundant regulators of hippocampal gene networks implicated in stress and mood disorders. This integrative molecular, behavioural, and endocrine framework identifies candidate regulatory mechanisms linking environmental adversity to hippocampal plasticity. Future studies involving targeted manipulation of miR-30e-5p and *Neurod1 in vivo*, coupled with direct assessment of neurogenesis, network function and behaviour, with larger sample sizes, could help determine their causal contribution to psychological stress-related cognitive outcomes.

## 9 Author contributions

The work was supervised by C.F. and S.T.. The study was conceptualised by C.F. and S.T. with support from A.M. who developed the hypothesis around miR-30e, *in silico* prediction of target genes and the object pattern separation paradigm to test for hippocampal function and S.T. with expertise in hippocampal neurogenesis and related cognition phenotypes. A.M., A.K. and C.F. performed the behavioural experiments. A.K. was the primary experimenter for behavioural scoring, with C.F. as expert for behaviour scoring used as secondary rater for inter-rater reliability measures. A.M. analysed the data with support from C.F. and S.T. . A.M. performed all gene expression and data analysis, methods, results, figures and supplementary data to produce this manuscript. A.M. and G.M. wrote the manuscript with input from the other co-authors

## 10 Funding

This work was supported by a UKRI Medical Research Council award reference 1667352.

### 11 Acknowledgements

The animal facility at Institute of Psychiatry, Psychology and Neuroscience (King’s Collect London) for supporting the animal routine husbandry and welfare.

