## Supplementary Information for "Social Isolation Alters Hippocampal miR-30e-5p Expression and Impairs Pattern Separation–Related Behaviour in Adult Mice"

**Supplementary Information (Supplementary Figures, Methods and Data)**

### Supplementary Figures

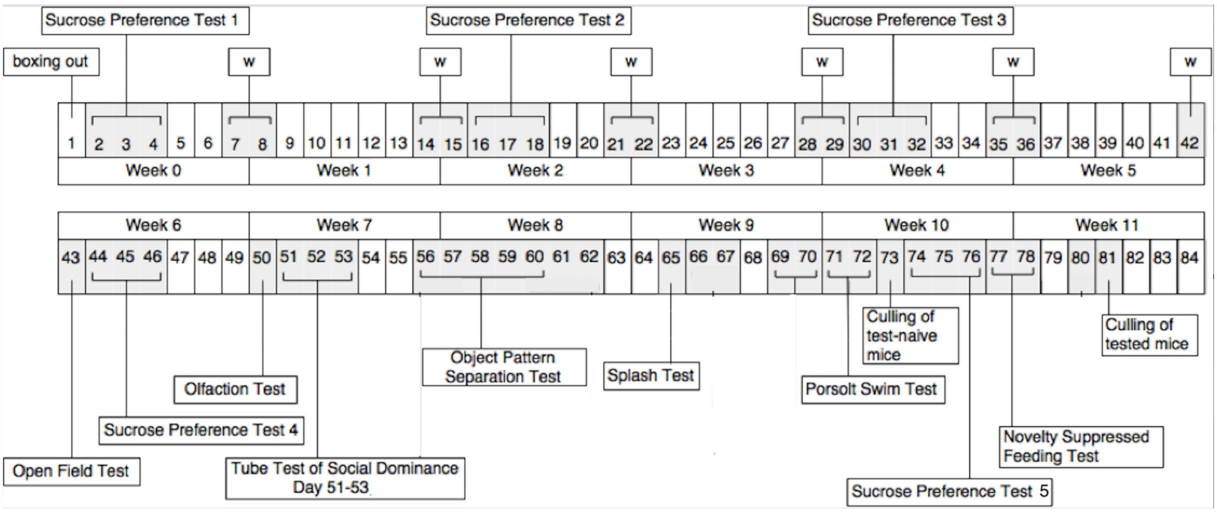


**Supplementary Figure 1 | Schematic timeline outlining the behavioural and cognitive testing schedule in the tested group of male BALB/cAnNCrl (BALB/c) mice.**

Twenty mice were subjected to behavioural and cognitive assessment (n=10 individually housed and n=10 pair housed mice) and results from individually housed and pair housed mice compared to determine whether social isolation by individual housing was associated with a significant anxiety/depression-like phenotype in male BALB/c mice. Testing was conducted in the order of least to most stressful to minimise impact of previous behavioural tasks confounding later test results. The experimenter responsible for measuring behavioural outcomes was blind to the housing group and hand scoring compared to an expert in behavioural analysis (inter-rater reliability intra-class correlation co-efficient > 0.80).

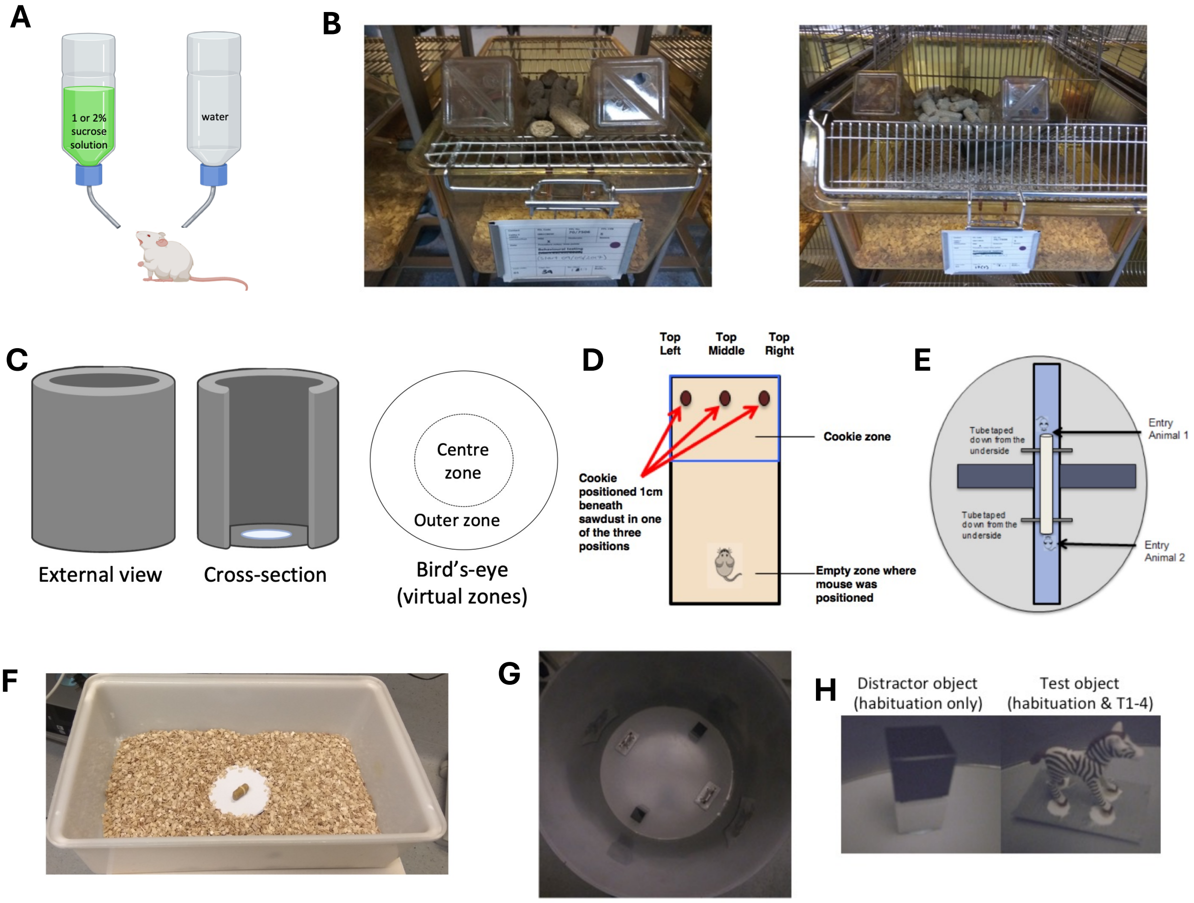


**Supplementary Figure 2 | Schematic diagrams and photographs of behavioural assessment and experimentation**

A) A schematic of the two-bottle test used to measure sucrose consumption over 48 hours (counterbalanced for side preference across the test period). (B) Photograph images of the two bottle set up for sucrose preference/consumption test for both (left) individually housed and (right) pair housed conditions. Sucrose solution (either 1% or 2% w/v) was prepared fresh and the bottles containing sucrose solution labelled with a small black circular sticker. C) Schematic outlining the open field test apparatus and virtual zone layout used by the tracking software (EthoVision) to record quantitative measures of open field avoidance. D) The buried cookie test is designed to test whether mice are able to detect and feed on a hidden cookie. E) A schematic **of** the tube test for social dominance. (F) A photograph image of the circular open field arena (40cm diameter) with objects positioned within the arena for habituation of the mice to exploration of the objects (image taken on day 1 of habituation). G) Photograph of the objects used for the habituation period. Distractor objects were used in habituation only to avoid a loss of interest in exploration of the test objects in advance of the test period, and to control for object preference. H) Photograph of test arena into which the mouse is placed after 24h food restriction (no access to food but water ad libitum).

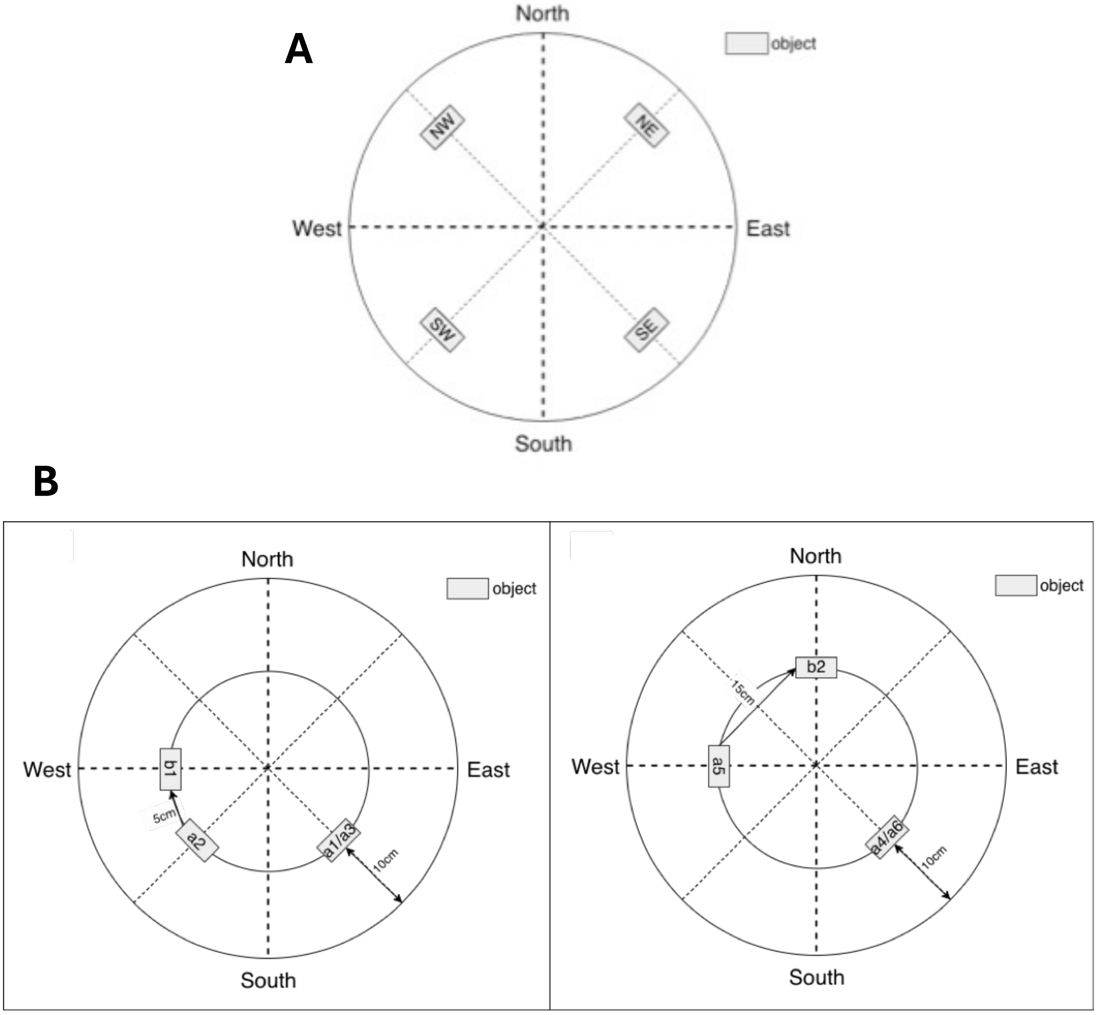


**Supplementary Figure 3 | Schematic diagrams of the arena for object pattern separation habituation and object pattern separation trials.**

A) Schematic indicating the virtual zones and objects positions (NW, NE, SW and SE) distanced 5 cm from the arena walls. Objects were counterbalanced between days such that distractor objects (left panel, Supplementary Figure 2G) in NE and SW day 1 and 3 of habituation, but NW and SE on day 2 and 4. Test objects (right panel, Supplementary Figure 2G) were in NW and SE positions on day 1 and 3, but NE and SW on day 2 and 4. B) Objects were positioned 10 cm from arena wall to avoid using an object location familiar to the mice from prior habituation. (B) Schematic of the arena in T1 to T2 (left panel), in which the test objects were positions 10 cm from arena wall in the SE and SW positioned in T1 and in T2 one object becomes displaced 5cm from starting SE position to occupy a position 10 cm from arena wall in the W direction whilst the static object remained in SW position. Schematic of the arena in T3 to T4 (right panel), in which the static test object remained positioned 10 cm from arena wall in the SW position and displace objected remained 10 cm from the arena wall in the W position.

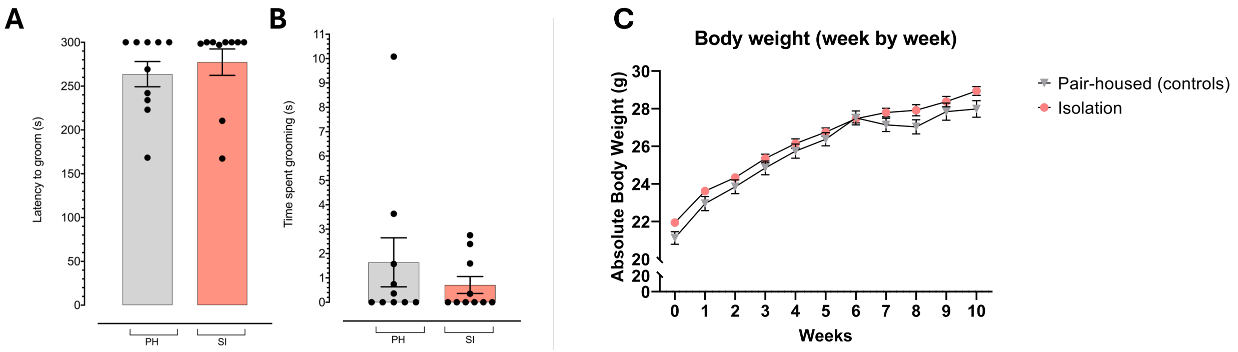


**Supplementary Figure 4 | Quantitative measures of grooming behaviour (Splash Test) and longitudinal body weight.**

The coat of each mouse was soiled with a sticky sucrose solution (10% w/v) in a novel cage environment in the Splash Test. No difference was observed in the (A) latency to groom and (B) time spent grooming in mice housed individually (SI, n = 10) compared to mice housed in pairs (PH, n= 10). Data graphed are mean ± SEM; each data point is representative of a measure from one animal. (C) In week 0, mice were assigned to either housing group matched for body weight. Body weight was recorded weekly (on Monday between 0900 and 1100) in the same order each week throughout the 10 weeks. No significant difference was observed in any week. Data graphed are mean ± SEM.

### Supplementary Methods

#### Coat state

The coat state of each animal was assessed weekly and a splash test conducted in week 9 to assess motivation to groom. Coat state is a measure mouse grooming, and validated for male BALB/c mice (Khemissi et al., 2014). Coat state was measured weekly using a scoring system as follows: the quality and condition of the animal's coat was scored according to a set of criteria which define the level of grooming where the total coat state score is the overall measure of grooming behaviour. Determination of the total coat state score (scale between 0 to 6) was achieved by calculating the cumulative score for seven individual body areas (head, neck, back, abdomen, forepaws and hindpaws where a score of 0 was given for well-groomed, 0.5 for moderately groomed and 1 for not groomed. Well-groomed was defined as smooth and shiny fur without any piloerection or patches. Moderately groomed was defined as having slight piloerection without any patches. Not groomed was defined as significant piloerection of the fur with patches and evidence of staining. Assessment was conducted weekly by the same experimenter.

#### Sucrose Preference Test

The Sucrose Preference Test (SPT) is a commonly used measure of preference for a sweet-tasting solution over water in mice and is suggested as a measure of anhedonia-like behaviour (Willner et al., 1987). Sucrose solution was prepared using room temperature drinking water and sucrose (1% w/v solution (in week 0, 2, 4 and 6) or 2% w/v solution (in week 10), S8501, Sigma, UK) (Supplementary Figure 2A and B). Sucrose concentration was increased in the final test to avoid habituation to sucrose from the previous tests in week 0-6) confounding the findings of the final test in week 10. A two-bottle test (Supplementary Figure 2B) was performed fortnightly during weeks 2, 4 and 6 and in week 10. Mice had *ad libitum* access to food and liquid over the test period where liquid was available from two identical bottles whereby one contained water and the other a sucrose solution (1% or 2% w/v). During the test period, consumption of sucrose solution and water was measured in grams (g) using weighing scales over 48 hours (divided into two, 24-hour test periods). Bottles were switched to alternate sides at the end of the first 24h test period until the end of the test 24h later to counterbalance for preference of drinking from a bottle on a particular side (i.e. in the first test period water was on the right and sucrose solution on the left side of the cage but in the second test period sucrose was on the left and water was on the right). Under normal conditions (i.e., not during the sucrose consumption test period), water was provided in two bottles to prevent development of preference for drinking from a bottle on one side by habituating animals to the two-bottle test environment.

To account for spillage, readings were taken from 6 spill cages and deducted from the total amounts consumed. Sucrose preference was calculated as the mass of sucrose consumed in the 48h test period (sum of amount consumed in first 24h period with amount consumed in second 24h period after each bottle's side location was swapped) as a percentage of the total liquid consumption (sum of water and sucrose consumed) per mouse:

$$\text{Sucrose preference (\%)} = (\Delta\text{mass}_{\text{sucrose 0-48h}} / \text{total consumption}_{\text{0-48h}}) \times 100$$

For pair-housed mice, consumption was assumed to be equal for the individuals within a pair. Data also were analysed combining consumption for the pair as a cage average and difference between cage means compared between groups and comparable results were obtained suggesting that analysis either way was not associated with different outcomes.

**Splash test**

The splash test is a measure of self-care by assaying motivation to groom upon contact with a sticky sucrose solution used to 'dirty' the animals' coat. The test assays motivation towards self-directed activities such as grooming (Yalcin et al.,

2005). The testing arena was an empty standard housing cage (test room light level 10 lux), with no sawdust or bedding. The mouse was sprayed with approximately 0.5ml of a 10% sucrose solution (Sigma, Gillingham, U.K.) onto their back at a range of 10cm. The mouse then was placed into the testing arena for 5 minutes. Grooming behaviour (latency to groom and time spent grooming) was assessed post-acquisition from a DVD recording.

**Open Field Test**

The Open Field Test (OFT) was used to assess anxiety-like behaviour (Gould, 2009). A circular (40 cm diameter x 40cm height) open field arena (Abplas, Mitcham, U.K.) was subdivided into virtual zones using movement-tracking software (EthoVision 3.1, Noldus, Wageningen, The Netherlands). The arena included a circular centre zone (20cm diameter) and an outer zone comprising of the outer ring of the arena (5cm width) (Supplementary Figure 2C). Each animal in the tested group underwent one trial (10-minute duration). At the start of each trial, the animal was placed in the outer zone of the arena and their movement tracked using a camera mounted vertically over the arena. The procedure room was maintained at constant lighting levels (10 lux), and the experimenter was absent from the room during testing.

**Olfaction Test**

The cookie test as a measure of olfactory function (Yang & Crawley, 2009) was conducted to control for anosmia, a potential confound for behavioural measures which involving feeding or olfaction (i.e. novelty suppressed feeding, sucrose preference and food consumption). A clean, standard mouse cage (32cm x 16cm x 14cm) containing a 3cm layer of fresh sawdust was set up as the test arena. Animals were placed in the arena for 5 minutes to habituate to the test environment. Animals were removed momentarily, whilst the experimenter hid a small cereal-based cookie (1cm diameter, Nestle Cookie Crisp®, UK) 1 cm below the surface of the sawdust (Supplementary Figure 2D). The animal then was returned to the arena and allowed a total exploration time of 5 minutes. The animal was placed at one end of the test arena in an empty zone. A qualitative outcome of YES was recorded for mice that were able to find the hidden cookie within the trial time. An outcome of NO was recorded for mice that did not find the cookie. Animals were scored as having no olfactory impairment if they were able to find and feed on the cookie within the 5 minutes test time. The procedure room was maintained at constant lighting levels (10 lux).

**Social Dominance Test (SDT)**

The tube test of social dominance (SDT) as previously described (Lindzey et al., 1961) was conducted to test for dominance within pair-housed mice as social hierarchy can affect behavioural outcomes (Beery & Kaufer, 2015). Social dominance was tested by using a 3.5cm diameter clear acrylic tube (Abplas, Mitcham, U.K) taped down with the entry points directed towards the open arms of an elevated plus maze (Abplas, Mitcham, U.K) in a procedure room maintained at constant lighting levels (10 lux), (Supplementary Figure 2E). Only pair-housed animals were tested, but individually house mice underwent a sham test in which they were placed into the test arena alone (at either of the two ends of the tube, counterbalanced within the group) as a sham test to control for exposure to the test arena. For the social dominance trial, cage mate pairs were places at opposite ends of the tube and released simultaneously. The natural tendency for animals is to travel forwards into the tube and attempt to traverse eventually encountering each other within the tube. In brief, cagemate pairs were positioned to enter either end of a clear acrylic tube of sufficient diameter (3.5 cm) to allow a single mouse to traverse. Upon meeting within the tube, the presence of a dominant cage mate was observed as the animal who proceeded forward within the tube thus forcing the other cage mate to reverse or turn around to exit the tube through the entrance in which they came. The animals underwent four trials in total, one two-minute trial per day for four days. A dominant cage mate was qualified as an animal that traversed the tube causing their submissive cage mate to rear backwards or turn to exit the tube via their entry point in at least 3 out of the 4 trials.

Animal would be recorded as the dominant cagemate if upon entering the tube and attempting to traverse from one side to another it proceeded forwards and ultimately (on at least 3 out of 4 tests counterbalanced to test alternate sides for each mouse) caused their submissive cagemate to regress towards their entry point to allow the dominant cagemate passage (Supplementary Figure 2E). Individually housed mice (n=10) underwent a sham test where they were permitted time to traverse the tube, alone. Test naïve mice intended for molecular analysis were not subjected to this test.

**Novelty-Suppressed Feeding (NSF) Test**

The NSF test examines hyponeophagia, the suppression of feeding due to exposure to a novel environment that has been used as a measure of anxiety in rodents. The NSF test arena consisted of a brightly lit (300 lux), white plastic cage (25x11x42cm) with a clear Perspex lid on top (not shown) to prevent escape. The cage was filled with sawdust (2cm deep), and a single food pellet was fixed (using a rubber band) to a plastic dish placed in the centre of the arena (Supplementary Figure 2F). After a period of food restriction (24h without access to food but *ad libitum* access to water) to induce a state of hunger in the mouse and increase food-motivated exploratory behaviour, mice were individually placed in the NSF test arena. The single pellet available to the mouse in the arena is the first time the mouse has had access to food for 24h. This test exploits rodents' motivation to feed (after food restriction) against the conflict of being in a novel, potentially threatening/anxiogenic arena. The mouse was permitted 5 minutes in the arena. Outcome measures were the latency to find the pellet and feed, and the amount of pellet eaten. Anxious mice would be predicted to take longer to enter the centre of the arena to feed and eat less of the pellet. The food pellet was weighed before and after the test to measure food consumption during the test. Homecage feeding also was assessed for 5 minutes immediately following their return to their homecage to assay motivation to eat in a familiar, non-threatening environment. To do this, mice were placed back into their homecage following the 5 minutes in the test arena and given a pre-weighed amount of food to eat. This amount of food was re-weighed, 5 mins post-trial and the difference between the weight before and after was taken as a measure of homecage feeding. The body weight of each mouse was recorded before food restriction and after homecage feeding (for welfare reasons) to measure effect of food restriction over a 24h period on weight loss.

**Porsolt Swim Test**

The Porsolt Swim Test measures the latency to, and time spent in, immobility when mice are placed in a cylinder filled with water (Porsolt et al., 2001). This immobile position is interpreted as the mouse giving up on escape and is an indicator of behavioural despair. On the test day, the mouse was placed into a tall, clear Perspex cylinder (49 cm high x 15 cm diameter) filled with room temperature water that was illuminated to 10 lux for 6 minutes and the experimenter exited the room. Immobility behaviour was scored post-acquisition. Following the test, the animal was transferred to their home cage to recover for 30 mins. A heat lamp was placed over half of the cage to provide warmth during recovery but also leaving a cooler area in the cage for the mouse to move to if it got too hot. The Porsolt swim test also was used as an acute stressor. Given that social isolation is a chronic mild stressor, an acute stressor was used and a blood sample taken before and after exposure to the acute stressor as a measure of HPA axis. Blood was taken 24h prior to, and 30 minutes after, the Porsolt swim test to minimize any impact of blood collection on the Porsolt swim test assessment.

**Object Pattern Separation Habituation**

Animals were habituated to the test arena in the presence of a pair of distractor objects (to avoid loss of interest in test objects) and a pair of test objects for 5 minutes per day over the course of 4 days. The distractor objects were black and white cubes (2cm x 2cm x 2cm) stacked one on top of the other (Abplas, Mitcham, U.K.) and the test objects were two black and white Playmobil© zebra figurines (3.5cm long x 4.5cm high) (Playmobil, Basildon, U.K.). The four objects (two distractors and two test) were placed in different quadrants of the arena (5cm from the arena wall) and, where appropriate, the same central orientation along either the NW-SE axis or NE-SW axis between the daily trials. The configuration of the objects on day 1 and 2 of habituation was reversed on day 3 and day 4 i.e. the blocks occupied the NW and SE positions while the zebra figurines occupied the NE and SW positions on day 1 and 2, and on day 3 and 4 the blocks occupied the NE and SW positions while the zebra figurines occupied the NW and SE (Supplementary Figure 3A). Animals were placed in the arena in the North position, directly at the meeting of the arena wall with the arena floor (facing the wall rather than into the arena). Exploration behaviour was not hand scored during habituation, but animals were tracked using Ethovision and the time spent inside each of the object quadrants was measured to determine a preference for a particular area or object in the arena. Data collected from animals that did not explore the central zone of the open field of the arena during habituation were omitted from the analysis. One single-housed mouse and one pair-housed mouse were excluded from the analysis at this stage since they did not enter the central zone during habituation. Objects were placed in the centre zone during testing; therefore lack of exploration of this zone of the area by a mouse could confound its exploration during the subsequent OPS trials.

**Object Pattern Separation (OPS) Task**

The OPS task was conducted 24 hours after the last habituation session for each mouse. All testing was conducted on the same day, with the same experimenters as during the habituation. The task was subdivided into four successive trials (4 mins per trial) separated by a 1-hour inter-trial interval. All mice underwent all four trials. Objects were positioned 10cm from, and parallel to, the arena wall to ensure the position of the test object was not in, or near to, the position of either the test object or distractor objects in during habituation. Trial 1 (T1) was an encoding trial for the purposes of acquiring a memory of the test context through experience prior to object displacement. In trials 2 (T2) and (T3), one test object in the SW position was radially displaced 5cm from the location in (T1) to occupy a position 10cm of from the arena wall but in the West position. Trial 4 (T4) was a final test trial in which the object was radially displaced a further 10cm to occupy a position in the North position having previously occupied a position in the West. Therefore, in T4, the object was displaced 10cm from the position occupied in T2 and T3 but 15cm radially from the position occupied in T1, the encoding trial (Supplementary Figure 3B). Object positions during the test were never overlapping with the object locations during habituation to exclude the possibility that animals were visiting the area where they expected to find an object. Exploratory behaviour was defined as the mouse directing their nose towards the object at a distance of no more than approximately 1cm and/or touching the object with the nose. Sitting near or on the object or lying/leaning near or on the object was not considered exploration. Exploratory behaviour was measured post-acquisition from a DVD recording, by the same experimenter with validation of inter-rater reliability. Data from animals that did not explore both objects for a minimum of 1 second were omitted from the analysis. Two single-housed and one pair-housed mice were omitted from the overall analysis due to a lack of object exploration in T1 which could confound results in subsequent trials since object exploration was insufficient in the first acquisition trial to assume the mice had acknowledged that two symmetrically placed objects were present in the arena. Objects were positioned 10 cm from arena wall to avoid using an object location familiar to the mice from prior habituation. (A) Schematic of the arena in T1 to T2, in which the test objects were positions 10 cm from arena wall in the SE and SW positioned in T1 and in T2 one object becomes displaced 5cm from starting SE position to occupy a position 10 cm from arena wall in the W direction whilst the static object remained in SW position. In T3 to T4, the static test object remained positioned 10 cm from arena wall in the SW position and displace object remained 10 cm from the arena wall in the W position. In

T4 the displaced object is moved again 15cm from W position to occupy a position 10 cm from arena wall in the N direction. An object discrimination index (*d*) was used as the outcome measure and was assessed for all trials (T1-4) where *d* = (*a2 – a1) / (a1 + a2)* in T1*, d = (b1 – a3) / (b1 + a3)* in T2*, d = (a5 – a4) / (a5 + a4)* in T3 and *d = (b2 – a6) / (b2 + a6)* in T4*.* A discrimination index (*d*) was calculated as the outcome measure for each animal in the trial. The total object exploration time (e.g., $Total_{T1}$ [s]), i.e., the time in seconds spent exploring both objects summed up, was calculated per trial to give a total object exploration time per trial for T1, T2, T3 and T4. The difference in exploration between objects was calculated (e.g., $\Delta_{T1}$ [s]), i.e., the time in seconds spent exploring the stationary object subtracted from the time spent exploring the displaced object. The discrimination index was calculated. Therefore, *d* is a number between -1 and 1, where -1 represents total time spent exploring the stationary object and 1 represents a total time spent exploring the displaced object and 0 represents equal time spent exploring both objects.

### Supplementary Data

**Supplementary Data 1 | Summary table of genomic targets of mouse NEUROD1 identified in the literature.**

| # TF | Target | Type | Reference (PMID) |
|---|---|---|---|
| Neurod1 | *Atoh1* | Activation | 20661473 |
| Neurod1 | *G6pc2* | Unknown | 12540293 &18753309 |
| Neurod1 | *Ins2* | Unknown | 16461554 |
| Neurod1 | *Insm2* | Activation | 21343251 |
| Neurod1 | *Mgat5b* | Activation | 21771782 |
| Neurod1 | *Nnat* | Unknown | 15793245 |
| Neurod1 | *Pax6* | Activation | 12962539 |
| Neurod1 | *Pdx1* | Unknown | 20448145 |
| Neurod1 | *Rnd2* | Activation | 23180754 |
| Neurod1 | *Sct* | Activation | 17875929 |
| Neurod1 | *St18* | Unknown | 23236509 |

**Supplementary Data 2 | Summary table of the Gene Ontology Biological Processes in which genomic targets of mouse NEUROD1 are enriched.**

| GOBP Term | GOBP Accession | [# of overlapped genes] | [P value] | [FDR] |
|---|---|---|---|---|
| type B pancreatic cell differentiation | GO:0003309 | 3 | 3.75E-07 | 8.77E-09 |
| glucose homeostasis | GO:0042593 | 4 | 9.19E-06 | 5.14E-07 |
| pancreatic A cell differentiation | GO:0003310 | 2 | 3.55E-05 | 3.19E-06 |
| regulation of gene expression | GO:0010468 | 4 | 0.000116793 | 1.64E-05 |
| brain development | GO:0007420 | 3 | 0.000278051 | 5.89E-05 |
| endocrine pancreas development | GO:0031018 | 2 | 0.000407509 | 0.000102897 |
| regulation of neuron differentiation | GO:0045664 | 2 | 0.000427871 | 0.000111092 |
| multicellular organism development | GO:0007275 | 5 | 0.000598005 | 0.000178339 |
| transcription, DNA-templated | GO:0006351 | 6 | 0.000856258 | 0.000294526 |
| response to glucose | GO:0009749 | 2 | 0.000952023 | 0.000339903 |
| positive regulation of insulin secretion | GO:0032024 | 2 | 0.001013686 | 0.000369532 |
| regulation of transcription, DNA-templated | GO:0006355 | 6 | 0.001305018 | 0.00052569 |
| glucose metabolic process | GO:0006006 | 2 | 0.001380728 | 0.000572779 |
| negative regulation of transcription from RNA polymerase II promoter | GO:0000122 | 4 | 0.001423108 | 0.000595024 |
| regulation of protein localization | GO:0032880 | 2 | 0.00148678 | 0.000630376 |
| cell differentiation | GO:0030154 | 4 | 0.001606032 | 0.000702087 |
| smoothened signalling pathway | GO:0007224 | 2 | 0.001617991 | 0.000711367 |
| negative regulation of protein phosphorylation | GO:0001933 | 2 | 0.002179617 | 0.001057466 |

| | | | | |
|---|---|---|---|---|
| negative regulation of cell proliferation | GO:0008285 | 3 | 0.002191238 | 0.001065769 |
| positive regulation of cell proliferation | GO:0008284 | 3 | 0.003604443 | 0.00205258 |
| neuron migration | GO:0001764 | 2 | 0.003838402 | 0.00222809 |
| axon guidance | GO:0007411 | 2 | 0.00456036 | 0.002763201 |
| transcription from RNA polymerase II promoter | GO:0006366 | 2 | 0.011282694 | 0.008465359 |
| positive regulation of transcription from RNA polymerase II promoter | GO:0045944 | 3 | 0.021090903 | 0.017455434 |
| nervous system development | GO:0007399 | 2 | 0.022211626 | 0.018531367 |
| regulation of transcription from RNA polymerase II promoter | GO:0006357 | 2 | 0.023518277 | 0.019803738 |
| negative regulation of transcription, DNA-templated | GO:0045892 | 2 | 0.035761714 | 0.031610946 |
| positive regulation of transcription, DNA-templated | GO:0045893 | 2 | 0.045782006 | 0.041472621 |

**Supplementary Data 3 | Summary table of the KEGG Pathways in which genomic targets of mouse NEUROD1 are enriched.**

| KEGG Term | KEGG Accession | [# of overlapped genes] | [P value] | [FDR] |
|---|---|---|---|---|
| Maturity onset diabetes of the young | mmu04950 | 3 | 3.77E-06 | 4.55E-07 |
| Type II diabetes mellitus | mmu04930 | 2 | 0.000974914 | 0.000384803 |
| Insulin secretion | mmu04911 | 2 | 0.002219107 | 0.001134658 |
| Insulin resistance | mmu04931 | 2 | 0.003261725 | 0.001879771 |
| AMPK signalling pathway | mmu04152 | 2 | 0.004110033 | 0.002527129 |
| FoxO signalling pathway | mmu04068 | 2 | 0.004365299 | 0.002723156 |
| Insulin signalling pathway | mmu04910 | 2 | 0.004717775 | 0.003009312 |
| PI3K-Akt signalling pathway | mmu04151 | 2 | 0.020076761 | 0.017109174 |
